# Reconstructing Prehistoric Waterscapes and Human Mobility during MIS 5 in North Africa

**DOI:** 10.64898/2026.07.19.739200

**Authors:** Solène Boisard, Colin D. Wren, Lucy Timbrell, Ariane Burke

**Affiliations:** Department of Anthropology, Université de Montréal, Montreal, Quebec, Canada; Department of Anthropology, University of Colorado Colorado Springs, Colorado Springs, CO, USA; Department of Archaeology, Classics and Egyptology, University of Liverpool, Liverpool, United Kingdom; Human Palaeosystems Group, Max Planck Institute for Geoanthropology, Jena, Germany

**Keywords:** Marine Isotope Stage 5 (MIS 5), North Africa, Human mobility, Paleohydrology, Optimal path analysis

## Abstract

The spatial organization of *Homo sapiens* populations in North Africa during Marine Isotope Stage 5 (MIS 5) remains a key question for understanding past demographic and cultural dynamics. Climatic fluctuations during this period produced alternating humid and arid phases, reshaping suitable habitats and influencing potential movement corridors. Previous studies have emphasized the importance of hydrographic networks for human dispersal, yet the extent to which climate-driven hydrological changes structured regional connectivity across MIS 5 substages remains poorly understood. This study integrates downscaled paleoclimate simulations, hydrological modelling, GIS-based optimal path analyses, and archaeological site distributions to reconstruct patterns of human mobility during MIS 5. We model palaeohydrographic networks for each MIS 5 substage and evaluate how precipitation variability influenced freshwater availability and landscape connectivity. Our results reveal spatio-temporal shifts in hydrographic networks and identify river corridors that likely facilitated movement within North Africa and the Saharan desert. These findings provide new insights into the role of waterscapes in structuring human mobility and contribute to broader discussions of Late Pleistocene human dispersal in Africa.

## 1. Introduction

The earliest known records of the emergence of *Homo sapiens* occurs in North Africa, dated 315 ka (Richter et al., 2017) and in East Africa, dated ca. 233□ka (Vidal et al., 2022). A series of dispersals out of Africa are thought to have occurred sporadically since 300ka, driven by climate conditions which facilitated human mobility in arid regions such as the Sahara (Timmermann and Friedrich, 2016; Scerri et al., 2018; Beyer et al., 2021).

Understanding the spatial organization of *Homo sapiens* during the last Interglacial, i.e., Marine Isotope Stage (MIS) 5 (∼130,000–70,000 years ago), is essential to reconstructing the demographic and cultural dynamics of our species. The greening of the Sahara during this period likely facilitated human movements both within and beyond the African continent (Blome et al., 2012; Larrasoaña et al., 2013; Scerri et al., 2019; Blanchet et al., 2021). Northwest Africa offers a crucial case study for examining how environmental conditions shaped human settlement patterns and connectivity. North Africa experienced increased humidity and the expansion of lakes and rivers, which may have supported human occupation of the Sahara and promoted contact between distant groups within North Africa and in sub-Saharan Africa (Blanchet et al., 2021; O’Mara et al., 2022). The archaeological record of the region is dominated by undated surface sites and sites whose chronology is typology-based, however (**Figure 1** - Scerri and Spinapolice, 2019; Boisard and Ben Arous, 2024).

**Figure 1.**
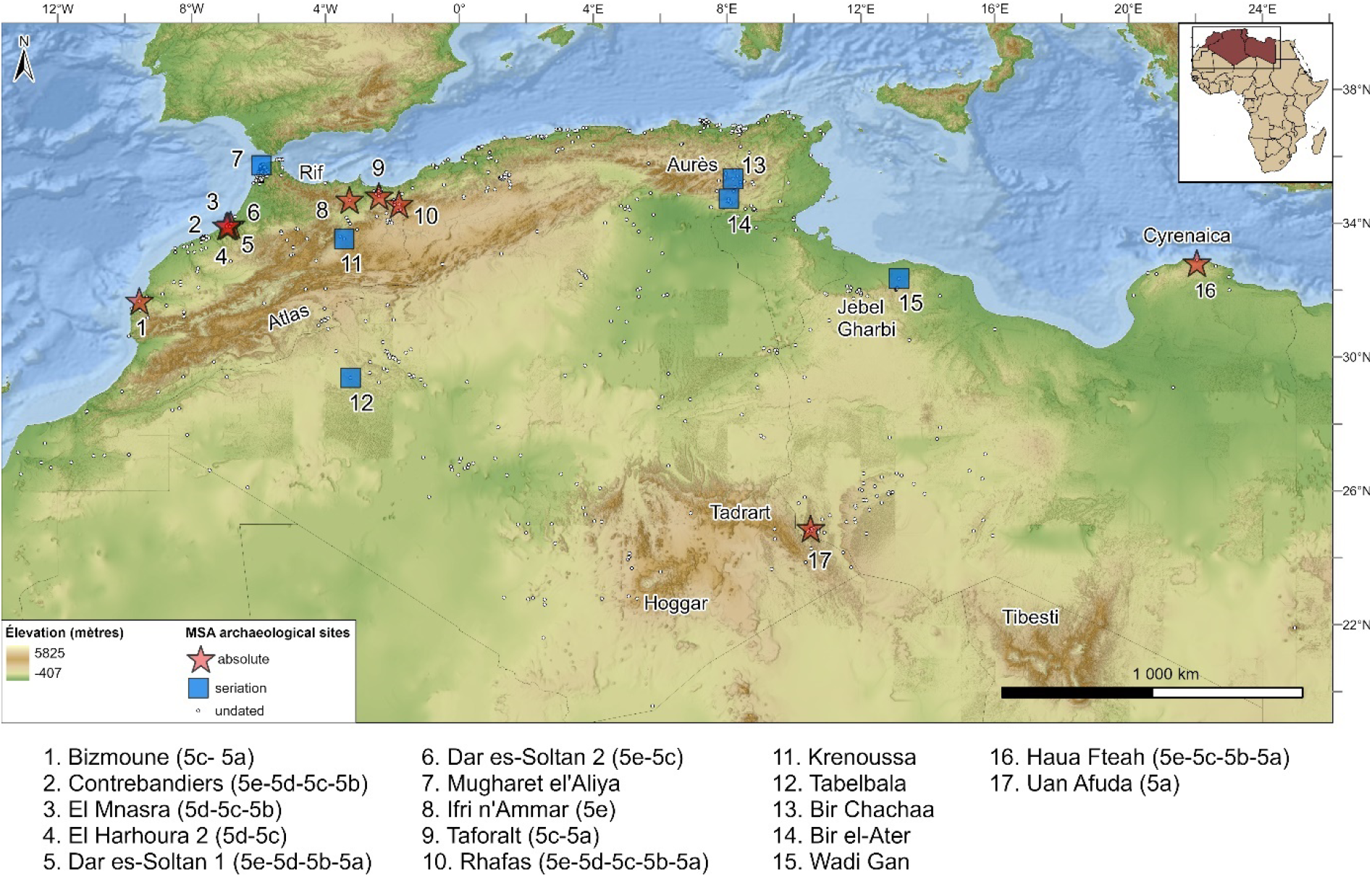
Distribution of MSA archaeological sites dated to MIS 5, with their chronological status based on Boisard and Ben Arous (2024), and identification of the main regions discussed in the text.

The Middle Stone Age (MSA) lithic industries of the region are marked by regional variability (Scerri and Spinapolice, 2019; Scerri and Will, 2023), but also by emerging symbolic behaviors such as the use of personal ornaments, pointing to expanding social networks and complex cultural dynamics (d’Errico et al., 2009; Barton and d’Errico, 2012; Hublin and McPherron, 2012; Van Peer, 2016). These developments suggest that human populations were engaged in social interactions and cultural transmission across groups (e.g., Scerri et al. 2018). Such patterns imply a degree of mobility and contact among populations that may have been facilitated – or constrained – by environmental factors. Lithic evidence suggests that North African populations were relatively well connected along the Atlantic littoral and the Jebel Gharbi, with shared technological traditions consistent with regional interaction, whereas Cyrenaica appears to have remained comparatively isolated (Scerri, 2013, 2013; Scerri, et al. 2014). The extent to which environmental factors influenced the spatial distribution of human populations, their mobility, and patterns of social interaction in the study region remains insufficiently explored, however (e.g, Scerri et al., 2014). This study addresses this gap by modeling suitable habitat and predicting patterns of mobility across different substages of MIS 5 in Northwest Africa. Using downscaled paleoclimate simulation output (Krapp et al., 2021), we identify areas of suitable habitat and reconstruct potential paleohydrographic networks. These networks inform optimal path models that predict likely routes between archaeological sites across the study region. By comparing site-to-site accessibility within each MIS 5 substage, we further evaluate how reconstructed hydrological networks structured regional connectivity and identify groups of sites that were more closely linked through time. Our goals are to (i) delineate core zones of human occupation, (ii) reconstruct potential movement corridors, (iii) evaluate how water availability structured connectivity among archaeological sites through time, and (iv) provide a framework for guiding future archaeological survey and excavation, particularly in regions where undated surface assemblages remain the primary source of evidence.

### 2. Regional and Temporal settings

The spatial extent of this study (18°N to 38°N and 15°W to 35°E) focusses on Northwest Africa, including present-day Morocco, Algeria, Tunisia, and Libya and a substantial portion of the Sahara Desert (**Figure 1**). All dated sites included in this study are located north of 25°N latitude, and as such, our primary focus is on the northern portion of the region. Northwest Africa is bordered by the Atlantic Ocean to the west and the Mediterranean Sea to the north, while the Sahara Desert dominates the interior, receiving less than 100 mm/year of precipitation on average today. The Atlas Mountains (∼4,165 m) play a crucial role in regional hydrology, generating orographic rainfall that supplies runoff to Saharan basins (Drake et al., 2022). Coastal ranges, including the Tell Atlas, Jebel Gharbi, and Jebel Akhdar, further contribute to localized runoff. Deeper inland, the Central Saharan massifs (e.g., Hoggar, Tibesti) act as secondary water sources, capturing moisture and sustaining local drainage systems (Cancellieri, 2021).

Paleohydrological studies indicate that past Saharan river networks were activated by the northward migration of the Intertropical Convergence Zone, which influenced monsoonal rainfall patterns across the Sahara (Larrasoaña et al., 2013; Pausata et al., 2020; Drake et al., 2022).

The MIS 5 chronology used in this study follows temporal intervals established in previous research (Kukla et al., 2002; Railsback et al., 2015; Creveling et al., 2017). The Interglacial period was marked by alternating humid and arid phases. MIS 5e (∼130-115 ka) marks the peak of humid conditions with widespread river activation and mega-lakes. These humid conditions may have persisted until ∼92 ka (e.g., Drake and Breeze, 2016), although MIS 5d (∼115-105 ka) is generally considered a more arid interval. Moderate humid episodes returned during MIS 5c (∼105-92 ka) and MIS 5a (∼85-70 ka), while MIS 5b (∼92-85 ka) represents another arid phase.

Numerous sources of evidence confirm the existence of several large lakes (>25,000 km²) in Northwest Africa during MIS 5 (Drake et al., 2022). Many of these low-lying basins, now covered by dune fields, were once filled with waterborne sediments during humid periods and later reworked by deflation in arid phases (see **SM**). Given the dating uncertainties associated with lake extent across distinct substages of MIS 5 (see Drake et al. 2022), this study incorporates the lake reconstructions by Drake et al., (2011) – a maximum lake extent model based on high humidity levels and interconnected basins – only as a comparative framework for evaluating the optimal pathways generated for each substage.

## 3. Materials and Methods

Climate data for this study was obtained using the *pastclim* package (Leanardi et al. 2023) in R (R Studio Team, 2020) to downscale and retrieve climate values from the Krapp et al. (2021) model dataset. Archaeological data preparation and spatial analyses were conducted in ArcGIS Pro 3.4 © ESRI. All code and datasets are publicly available online (see **Data availability**).

### 3.1. Archaeological data

The archaeological dataset is derived from a published database documenting MSA and LSA sites in Northwest Africa, including both dated and undated sites (Boisard and Ben Arous, 2024). For this study, we selected MSA occupations dated to 130–70 ka, classified as « contextual » in the original database – meaning their dates are directly associated with cultural materials, features (e.g., hearths), or human remains. The dataset includes 11 directly dated sites and 6 additional sites assigned to MIS 5 by seriation based on typological attributes (Scerri, 2013; Scerri et al. 2014). The attribution of archaeological site occupations to MIS 5 sub-stages followed a previously published protocol (Blinkhorn et al., 2022; Boisard et al., 2025). For each occupation, upper and lower chronological boundaries were established by applying ±1 standard deviation to the available age estimates. Where multiple dates were available, the oldest age (+1 s.d.) defined the upper boundary of occupation, whereas the youngest age (-1 s.d.) defined the lower boundary. The mean of all available ages was used as the representative occupation age for palaeoclimatic data extraction. This approach allowed each occupation to be assigned to the appropriate MIS 5 sub-stage(s). The final dataset comprises 17 archaeological sites and 61 dated occupations (**Table 1**).

**Table 1.** Northwest African dated archaeological sites and number of dated occupation(s) for each MIS 5 sub-stages (age between brackets). Sites dated by seriation not included (see Figure 1).

| Site | Acronym | 5e<br>(130-115) | 5d<br>(115-105) | 5c<br>(105-92) | 5b<br>(92-85) | 5a<br>(85-70) | Total |
| --- | --- | --- | --- | --- | --- | --- | --- |
| Bizmoune | BIZ |  |  | 2 |  | 1 | 3 |
| Contrebandiers | CTB | 2 | 5 | 6 | 2 |  | 15 |
| El Mnasra | EM |  | 1 | 3 | 2 |  | 6 |
| El Harhoura 2 | EH 2 |  | 1 | 1 |  |  | 2 |
| Dar es-Soltan 1 | DES 1 | 2 | 3 |  | 2 | 2 | 9 |
| Dar es-Soltan 2 | DES 2 | 1 |  | 1 |  |  | 2 |
| Ifri n'Ammar | IFR |  |  |  |  |  | 1 |
| Taforalt | TAF |  |  | 2 |  | 1 | 3 |
| Rhafas | RA | 1 | 1 | 1 | 2 | 1 | 6 |
| Haua Fteah | HF | 7 |  | 2 | 1 | 3 | 13 |
| Uan Afuda | UA | 1 |  |  |  | 1 | 1 |
| **Total** |  | **14** | **11** | **18** | **9** | **9** | **61** |

### 3.2. Climate and environmental data

Climate values used in this research were extracted from a delta-downscaled version of the Krapp et al. (2021) output, a bias-corrected model time series based on the HadCM3 General Circulation Model (GCM) (Valdes et al., 2017), covering the last 800,000 years in 1,000-year time intervals Following the delta-downscaling method presented in Timbrell et al. (2024), climate data from Krapp et al. (2021) were downscaled to a 5×5 arc-minute resolution (∼9.28 km). Briefly, downscaling is performed one monthly variable at a time (e.g. January temperature) by taking the original model output (at 30x30 arc-minute resolution) and a corresponding set of high-resolution modern simulations (WorldClim2; Fick and Hijmans, 2017), as well as the ETOPO2022 global relief model at 15x15 arc-minute resolution, resampled to 5x5 arc-minute resolution (NOAA National Centers for Environmental Information, 2022). Through integrating both bathymetric and topographic values for masking sea level changes, a delta raster is computed, adding the difference between past and present-day simulated climate to present-day observed climate (Timbrell et al. 2024). The resulting high-resolution monthly datasets are then used within *pastclim* (Leonardi et al. 2023) to recompute the 17 bioclimatic variables available in the original dataset (Krapp et al. 2021), from which we extracted annual temperature (bio01) and total annual precipitation (bio12).

For hydrological modeling, we selected bio12 (downscaled) and calculated the mean for each MIS 5 substage, resulting in one averaged precipitation raster per period (Table 2). Topographical variables were derived from the global SRTM 3 arc-second (∼90 m) Digital Elevation Model (DEM) (USGS, 2017). The Digital Elevation Model (DEM) was used to calculate slope using ArcGIS Pro’s Slope tool (in degrees by default) (see **SM**). To model travel cost, we applied Tobler’s Hiking Function (Tobler, 1993), which estimates walking speed (km/h) based on terrain steepness. Since the function requires mathematical slope (rise/run) rather than degrees or percent as input (Herzog 2020; White and Barber 2012), we converted the slope values to radians and applied the tangent function in the Raster Calculator (see **SM**). This transformation ensured that Tobler’s formula was applied correctly, producing a raster of walking speed that was then inverted to generate a cost surface, where flat terrain corresponds to lower cost and steep slopes to higher cost.

**Table 2.** Climate envelopes of annual temperature and precipitation for each MIS 5 substage used in the climate-archaeology models.

| MIS | Min temp.<br>(°C) | Max temp.<br>(°C) | Min prec.<br>(mm) | Max prec.<br>(mm) |
| --- | --- | --- | --- | --- |
| 5e | 13.7 | 18.1 | 326 | 473 |
| 5d | 13.7 | 17.3 | 472 | 485 |
| 5c | 13.7 | 18.1 | 314 | 485 |
| 5b | 13.7 | 18.1 | 373 | 485 |
| 5a | 15 | 23.8 | 19 | 485 |

### 3.3. Climate and hydrological models

#### 3.3.1. Climate-archaeology models

We applied the approach described in Boisard et al. (2025) to produce climate envelope models based on the location of archaeological sites and paleoclimate simulations. To this end, each dated archaeological layer was assigned its closest 1000-year timeslice and mean annual temperature (bio01) and total annual precipitation (bio12) values were extracted from the climate data. We then used the minimum and maximum temperature and precipitation values recorded at archaeological sites to define suitability ranges for each MIS 5 substage (**Table 2**). We conducted a frequency analysis of these climate envelopes to determine how often each grid cell falls within the defined temperature and precipitation ranges for each substage, as well as within the combined temperature and precipitation ranges. Based on these results, we calculated suitability expressed as percentages – where 100% indicates that a cell always meets the defined climate conditions throughout the substage, and lower percentages reflect decreasing frequency of suitability. We then projected these suitability scores spatially to generate maps of potential habitat for each MIS 5 substage. Grid cells with higher suitability values represent areas where climatic conditions repeatedly aligned with the reconstructed climate preferences of human populations adapted to this region’s climate states. We interpreted these values in terms of landscape types using the Holdridge (1967) life zone classification system. As such, the resulting maps highlight zones of potential ecological stability that may have supported sustained or recurrent human occupation.

#### 3.3.2. Hydrological models

To illustrate latitudinal variability in precipitation within the study area, we divided the region into northern and southern domains for visualization purposes only, using 25°N as a reference line (ssee **Figure 4**). This threshold corresponds to the northernmost reach of the West African monsoon during exceptionally humid periods (Gasse, 2000). This arbitral boundary was not used in our hydrological modeling and te hydrographic analyses were conducted across the full spatial domain of the study, which includes all northward-draining watersheds.

Previous hydrological reconstructions in Saharan regions have primarily relied on visual interpretation of satellite imagery (e.g., Drake et al. 2011) or on probabilistic simulations at a single time slice over localized areas (e.g., Coulthard et al., 2013). Here, we develop a GIS-based hydrological model – adapted from Ames (2014) – that integrates paleoprecipitation data from downscaled climate simulations (Krapp et al. 2021) to predict surface water availability across the MIS 5 substages (**Figure 2**). We used a DEM (250 meters) – resampled from the original 90-meter SRTM dataset (USGS) – and applied the Flow Direction tool in ArcGIS Pro (Spatial Analyst toolbox) with the D8 algorithm to determine the direction of flow of each cell, assigning drainage to the steepest downslope neighbor. Precipitation rasters were first resampled to match the DEM resolution and then used to weight the flow accumulation algorithm for each substage. The Flow Accumulation tool calculates the number of upstream cells contributing to flow at each location with the original volume per cell being derived from the annual precipitation values (mm/year). We tested three flow accumulation thresholds for MIS 5e: 1,000,000; 5,000,000; and 10,000,000 mm of accumulated precipitation to explore variations in drainage density and network extent. While the model does not explicitly account for evaporation or infiltration rates, applying higher thresholds serves as a proxy for conditions favoring the development of more persistent surface flows. The three test thresholds were applied in an exploratory analysis without prior calibration (**SM – Figure 12**). The resulting stream networks were then compared with existing hydrographic reconstructions, including Drake et al. (2011) and HydroRIVERS (Lehner and Grill, 2013), to identify similarities and differences in major drainage patterns and stream hierarchy (**SM – Figure 11**). Based on the observed results, a threshold of 10,000,000 mm was adopted for modeling all the MIS 5 substages to ensure consistency in the representation of major drainage systems. While our model also approximates the maximum extent of large endorheic basins, it does not account for precipitation-weighted flow into these basins (i.e., how full they might have been) so we do not use them within the analysis below.

**Figure 2.**
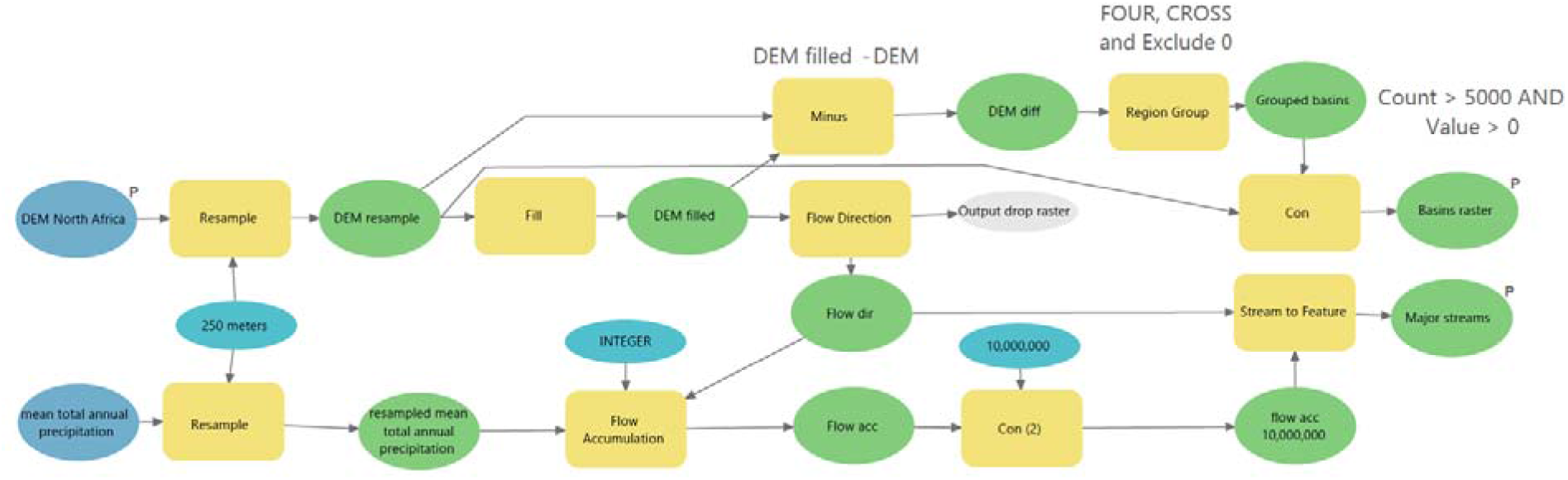
ModelBuilder workflow used for hydrological modeling (ArcGIS Pro 3.4).

### 3.4. Optimal Path models

We conducted an optimal path analysis using the Optimal Path as Line tool in ArcGIS Pro (Spatial Analyst toolbox) to identify potential human movement for each MIS 5 substage, integrating three key environmental factors as cost constraints: travel time (based on Tobler’s hiking function), precipitation, and Euclidean distance to modelled watercourses. Travel time represents the energetic cost of traversing the landscape, while annual precipitation serves as a proxy for broader environmental suitability, assuming that wetter areas supported greater vegetation productivity and terrestrial resource availability. Distance to watercourses represents direct access to freshwater resources, which are assumed to have been a primary constraint on human mobility in arid environments. Each factor was converted into a normalized cost raster and combined using the Weighted Sum tool in ArcGIS Pro. We tested four weighting scenarios to evaluate model sensitivity: water-weighted, slope-weighted, precipitation-weighted, and a balanced model. Weights represent the relative influence of each factor in the combined cost surface calculation (**Table 3**). The precipitation-weighted and slope-weighted models are presented in the **SM – Figure 13** for comparison but were not considered further because they overemphasize a single environmental variable while underrepresenting the importance of direct access to freshwater, a core assumption of this study (**Table 3**). The analysis assumes that individuals had sufficient landscape knowledge to select routes minimizing travel effort, consistent with least-cost path approaches (Herzog, 2013). Using the resulting weighted cost surfaces, each archaeological site was designated as an origin point in turn, and optimal paths were calculated to all other sites (**Figure 3**). This process generated a network of optimal paths for each substage, which were then visualized using the Line Density tool to highlight areas of concentrated movement potential (e.g., Paquin 2024).

**Figure 3.**
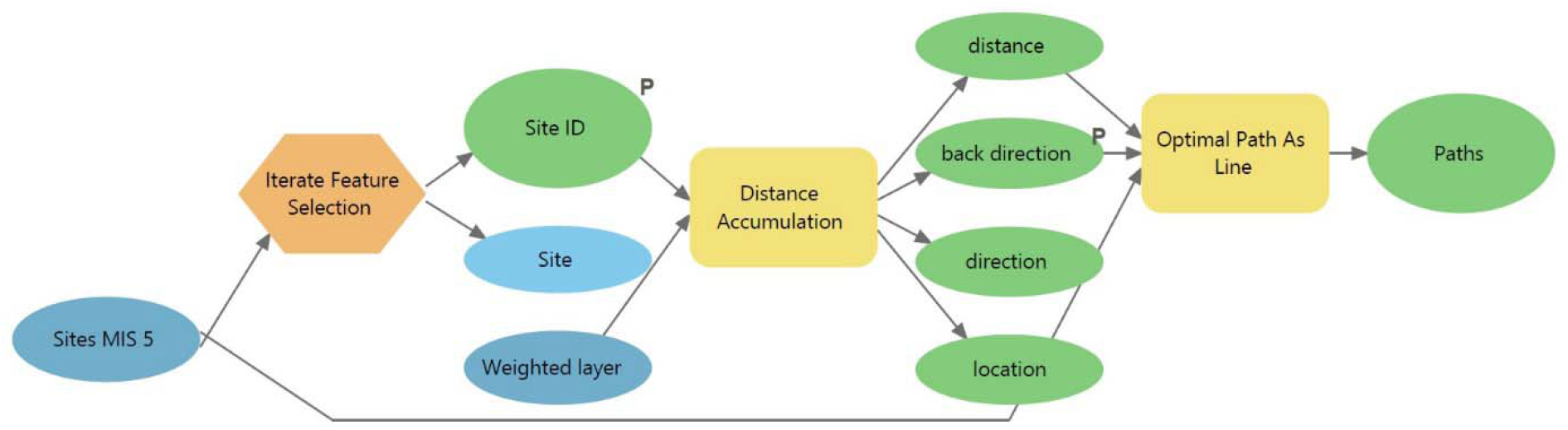
ModelBuilder workflow used in the Optimal Path modeling (ArcGIS Pro 3.4).

**Table 3.** Weighting scenarios used in the optimal path analysis.

| Scenario | Distance to Water | Slope | Precipitation | Reason for Inclusion/Exclusion |
| --- | --- | --- | --- | --- |
| Water | 0.90 | 0.05 | 0.05 | Included – Highlights influence of reconstructed hydrology on movement. |
| Slope | 0.05 | 0.90 | 0.05 | Excluded – Unrealistic detours paths in flat Saharan regions. |
| Precipitation | 0.05 | 0.05 | 0.90 | Excluded – Overemphasizes coastal routes due to low simulated rainfall inland. |
| Balanced | 0.34 | 0.33 | 0.33 | Included – Offers integrated perspective combining all environmental constraints. |

We then conducted a co-similarity analysis using the water-weighted model, which incorporates reconstructed hydrographic networks to estimate site-to-site accessibility and evaluate the influence of water availability on potential human movements during each MIS 5 sub-stage. Cumulative travel costs between each pair of sites were obtained from the cost surfaces generated with the Optimal Path tool in ArcGIS Pro. The resulting pairwise cost matrices were processed in R and transformed into co-similarity scores ranging from 0 (least accessible) to 1 (most accessible). This normalization emphasizes relative rather than absolute accessibility, enabling direct comparison across substages. Similar pairwise cost-based approaches have been used to investigate regional mobility and site clustering in archaeological landscapes (Herzog, 2013). The resulting matrices were visualized as heatmaps, revealing spatiotemporal clusters of relatively well-connected sites and highlighting how hydrological conditions structured regional connectivity throughout MIS 5.

## 4. Results

### 4.1. Hydroclimate regimes

Downscaled palaeoclimate data show that while there is substantial temporal variability in precipitation levels across MIS 5, most of that variability occurs in the southern part of the study area (**Figure 4**). Northern regions maintained relatively stable annual precipitation throughout the period, whereas the southern domain experienced greater variability, with increased precipitation during MIS 5e, end of 5d and 5a, indicative of intensified monsoonal influence. Hydrological reconstructions suggest the presence of perennial river systems, particularly in northern regions and major Saharan basins (**Figure 5**). The spatial extent of fluvial activity varied across substages in keeping with fluctuating precipitation rates, with MIS 5e showing the most extensive network of streams and lakes. This network contracted during dr er phases, especially MIS 5d and MIS 5b.

**Figure 4.**
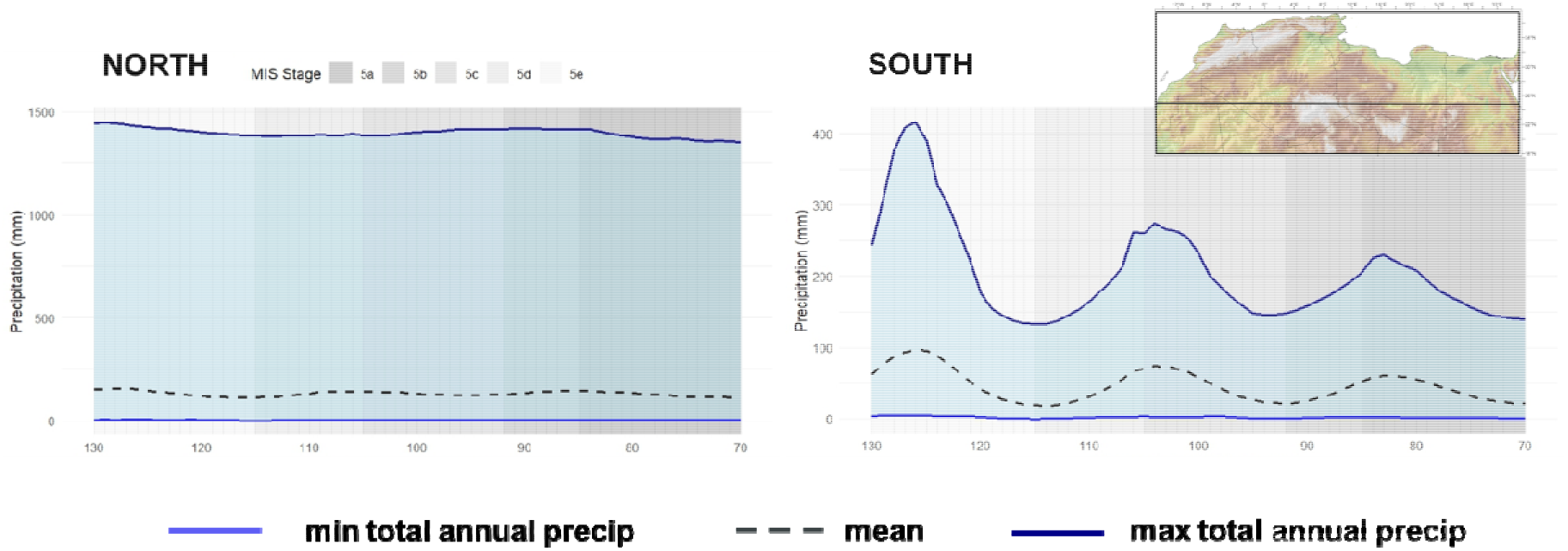
Distribution of total annual precipitation during MIS 5 based in northern (25-38 °N) and southern (18-25°N) areas for the study region. Values are erived from downscaled palaeoclimate simulations (Krapp et al. 2021). For each region and time slice, the dashed black line represents the mean total annual precipitation across all grid cells, the thin blue line indicates the minimum, and the thick dark blue line shows the maximum. MIS 5 substages (5e–5a) are shown as shaded background intervals. The inset map indicates the spatial extent of the northern and southern sectors used for averaging.

**Figure 5.**
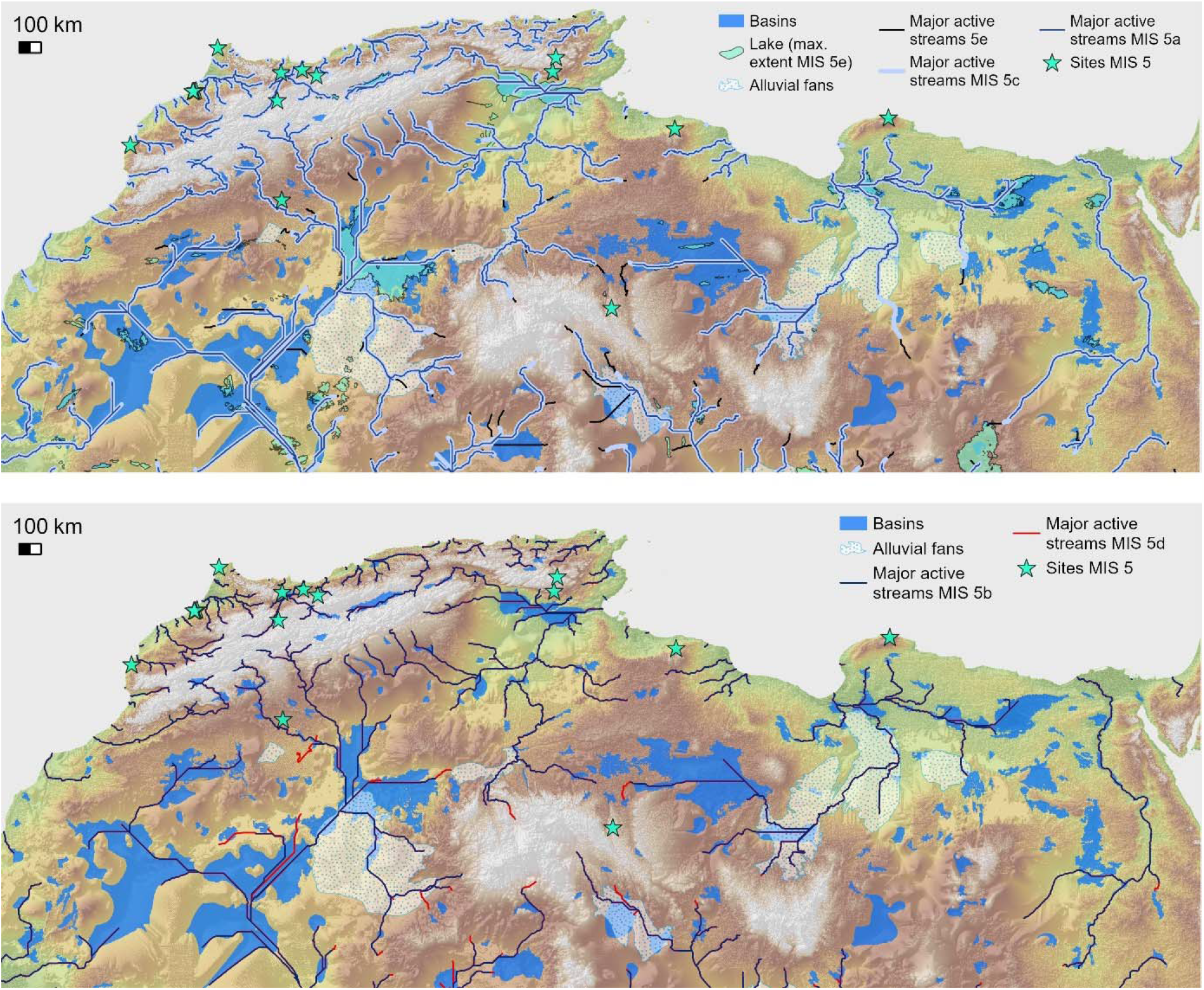
Simulated major streams for the wetter MIS 5e, 5c, and 5a (top) and drier MIS 5d and 5b (bottom), displayed with MIS 5 archaeological sites. Hydrographic networks were generated using a flow accumulation threshold of 10,000,000 mm and overlain with lake extents and alluvial fans from Drake et al. (2011). Active water systems from different substages partially overlap, highlighting both spatial continuities and shifts in surface water availability.

The MIS 5e paleohydrographic reconstruction shares broad similarities with both the Saharan river network proposed by Drake et al. (2011) and the modern high-order rivers mapped in HydroRIVERS (Lehner and Grill, 2013 – see **SM – Figure 11**). Major systems such as the Chéliff, Moulouya, Drâa, and Irharhar rivers appear in all datasets, as does the Nile, confirming that these water streams were reactivated during MIS 5. Our model also reveals a denser and more continuous fluvial network across interior regions – where the Saoura, Daoura, Fezzan, and Sahabi rivers flow. Several paleochannels, particularly in the Ahnet-Mouydir plateau and southwestern Libya, emerge as active under MIS 5e climatic conditions in our simulation but are absent in the modern datasets, highlighting the capacity of our model to identify hydrologically plausible but topographically subtle drainages. Conversely, certain rivers mapped with high Strahler orders (5-6) in HydroRIVERS are not reactivated in our simulation, likely due to insufficient simulated rainfall in Saharan areas.

### 4.2. Climate-archaeology models

Climate-archaeology models highlight substantial variation in the spatial extent of potentially habitable areas across MIS 5 substages (Figure 6). Climate data extracted from dated archaeological layers, detailed in the Appendix, show relatively stable values across substages in Atlantic and Mediterranean regions. Most temperature values range between 13.7°C and 18.2°C, and precipitation between 315 and 485 mm. These values define the climatic envelope used in the climate-archaeology models. MIS 5e, 5d, 5c, 5b exhibit similar envelopes, with suitable conditions present along the Atlantic littoral, the Mediterranean coast, and in Cyrenaica. MIS 5d, however, shows a marked spatial contraction of suitable areas, particularly along the western coast. This contraction results from the narrow climatic envelope reconstructed for this substage, especially with respect to precipitation (472-485 mm). Because the climate-envelope model is derived from the range of values recorded at archaeological sites, the relatively small number of dated MIS 5d occupations further restricts the range of suitable conditions represented.

**Figure 6.**
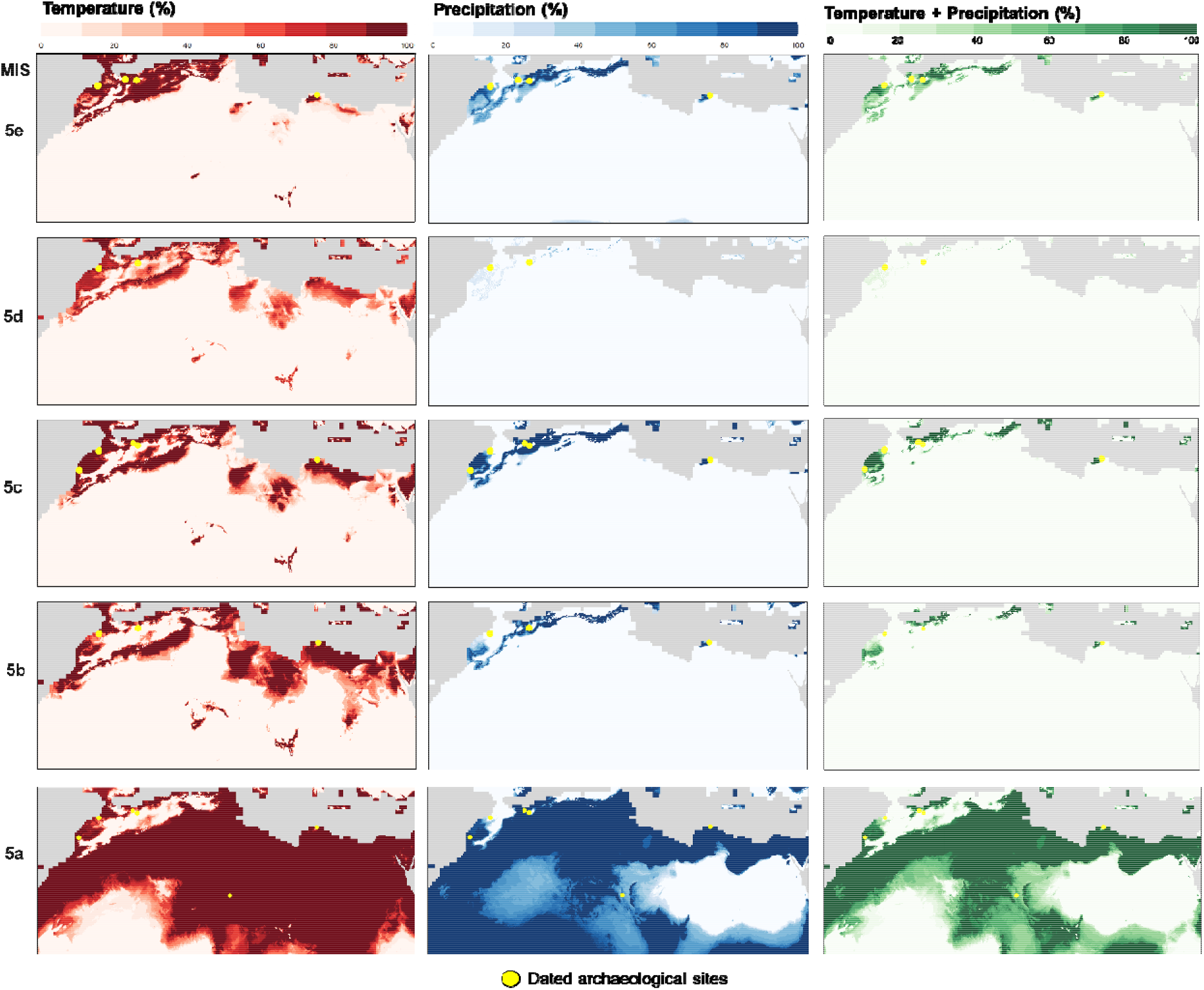
Climate-archaeology models for MIS 5 substages. Color gradients reflect the percentage of grid cells falling within the climate envelopes of temperature, precipitation and combined temperature-precipitation values documented at dated archaeological sites.

Although less pronounced than in MIS 5d, MIS 5b also exhibits a restricted spatial extent of suitable areas. Unlike the temperature and combined temperature-precipitation models, the precipitation model identifies MIS 5a as the period with the broadest extent of climatically suitable areas across the Sahara (**Figure 4**). This pattern contrasts with palaeoclimatic evidence indicating that MIS 5e represented the warmest and wettest interval of MIS 5. Rather than reflecting genuinely more favourable environmental conditions during MIS 5a, this result highlights a limitation of climate-envelope approaches based solely on temperature and precipitation. By ignoring the spatial distribution of freshwater resources, these models overestimate potential habitability across arid regions. This limitation underscores the value of incorporating reconstructed hydrographic networks, which provide a more realistic representation of landscape accessibility and better explain the observed distribution of archaeological sites (**Figure 1**).

### 4.3. Optimal paths models

The spatial configuration of optimal paths generated by the water-weighted and balanced models varies across MIS 5 substages (**Figure 7**). Both identify ecologically plausible movement corridors that closely follow reconstructed hydrographic networks, although the balanced model allows a greater influence of topography. For comparison, the precipitation-weighted and slope-priority models are presented in the Supplementary Materials (**SM – Figure 13**). These models overemphasize either precipitation or topographic gradients, producing pathways that are less consistent with reconstructed drainage systems and are therefore not discussed further here. In particular, slope-based models often generate long, linear routes that cross environmentally implausible areas, highlighting the limitations of relying on topography alone to infer potential movement corridors.

**Figure 7.**
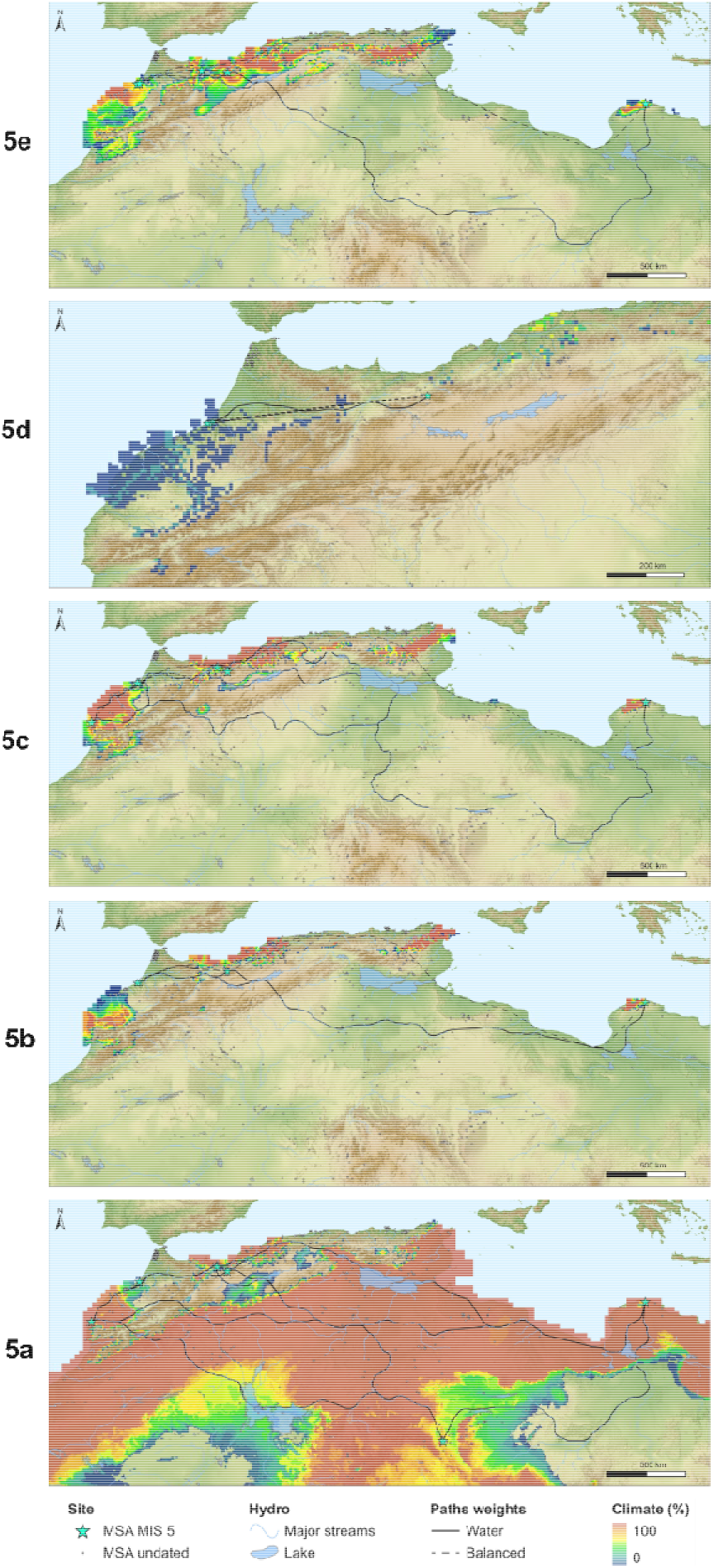
Maps combining climate-archaeology models and optimal paths (water and balanced) for each MIS 5 substage.

Across MIS 5e, MIS 5c, and MIS 5a, the water-weighted and balanced models consistently identify corridors linking northern and central Saharan archaeological sites through palaeodrainage systems. Uan Afuda repeatedly lies along these reconstructed routes, despite being dated only to MIS 5a, suggesting that the Hoggar region remained connected to broader movement networks during multiple MIS 5 substages. These connectivity patterns are consistent with previous hydrographic reconstructions proposing humid dispersal routes across the Sahara during MIS 5 (Osborne et al., 2008). Although Osborne et al. focused farther east, our results extend this interpretation westward by demonstrating that reconstructed drainage systems in the central Sahara also formed plausible pathways for human movement.

Pair-wise cumulative travel cost values (**Figure 8**), derived from optimal paths calculated in R using a water-weighted cost surface (see **Methods**), reveal that most sites exhibit similaraccessibility profiles when occupied. This suggests generally high inter-site connectivity across MIS 5. Coastal sites such as Dar es-Soltan 1 (DS1), Dar es-Soltan 2 (DS2), Ifri n’Ammar (IFR), Contrebandiers (CT), and Rhafas (RH) are consistently well integrated into an inter-accessible mobility network, especially during MIS 5e and 5d. Some sites, however, show more distinct patterns. Rhafas (RH), while usually central, shows reduced co-similarity in MIS 5a, likely due to shifts in paths orientation. Uan Afuda (UA), only occupied in MIS 5a, also appears marginal, situated along southern pathways that remain peripheral within the reconstructed network. Other sites, such as Taforalt (TAF) and Bizmoune (BIZ), vary in their connectivity depending on the substage.

**Figure 8.**
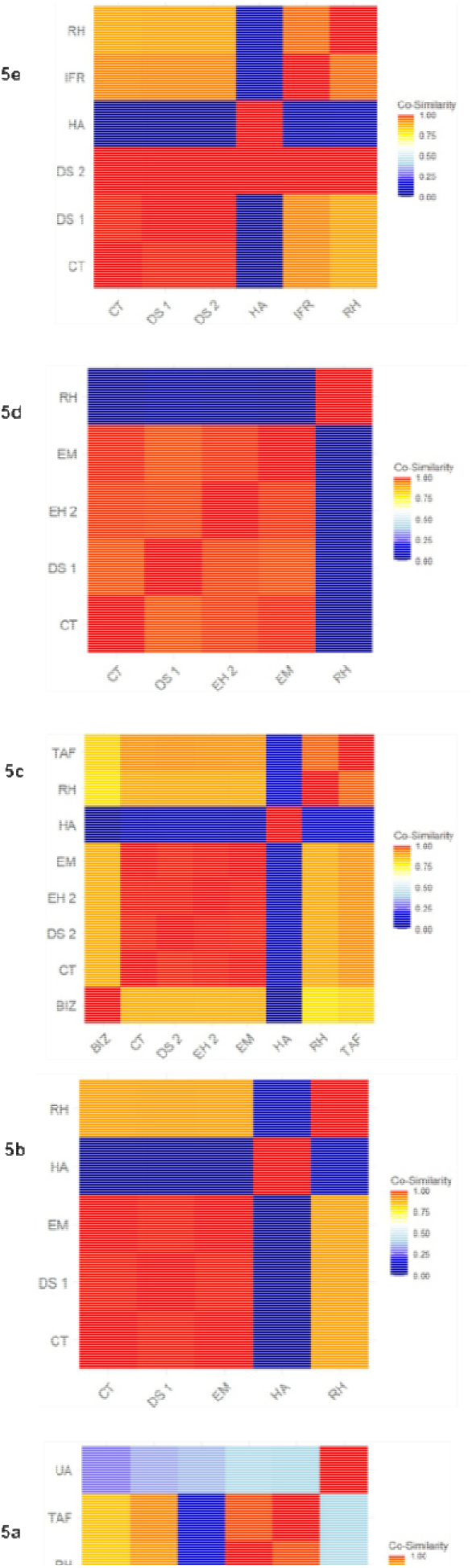
Co-similarity matrices calculated based on the cumulative cost of paths between each site for each substages

The general model combining all MIS 5 sites (dated and seriated) reveals a strong Atlantic cluster (**Figure 9**). Sites such as TAB, MUG, KRE, and CHA, which are dated by seriation to MIS 5 but cannot be assigned to a specific substage, show intermediate co-similarity with both coastal and inland sites. TAB stands out for its transitional position – linked to both northern and southern pathways, but less connected to eastern sites like HA or UA. Wadi Gan (WG) is well integrated with both Atlantic and Saharan nodes, especially UA, while HA remains consistently isolated, with a distinctly high mobility cost profile across the network.

**Figure 9.**
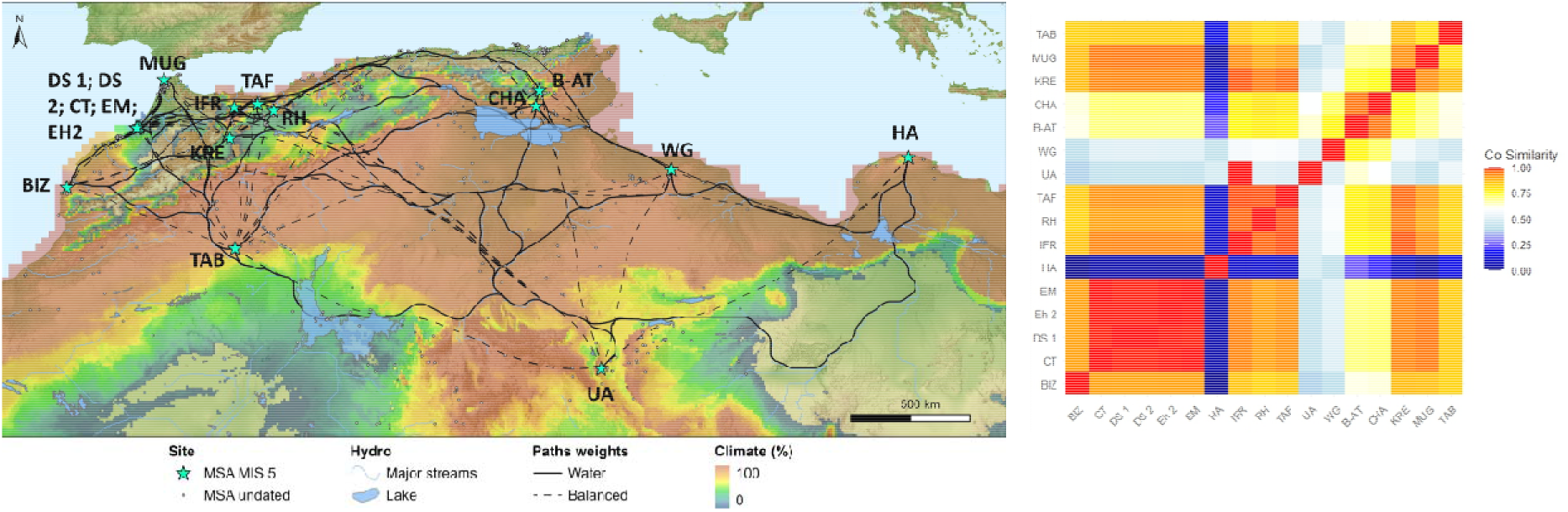
*Left*. MIS 5 General model combining climate-archaeology model f r the entire period and optimal paths of all dated (radiometric and seriations) MSA sites. *Right*: Co-similarity matrices calculated based on the cumulative cost of paths between each site for each substages.

## 5. Discussion

This study investigates the distribution of waterscapes and potential patterns of mobility in Northwest Africa during MIS 5 by integrating archaeological data, paleoclimate simulations, and hydrological modeling. We generated hydrographic reconstructions covering both Saharan and coastal river systems, revealing new potential corridors for human dispersal across Northwestern Africa and into the Central Sahara that may have played an important role in continental-scale human dispersals, both within and beyond Africa (Osborne et al., 2008; Stewart and Stringer, 2012; Scerri et al., 2019; Beyer et al., 2021). We combined climate-envelope modelling with hydrological reconstructions and optimal path analyses to examine how environmental factors influenced human mobility across MIS 5 substages.

Before interpreting our results further, it is important to recognise the limitations of this study. First, although we used spatially downscaled climate simulations, some coverage gaps are already present in the original dataset (Krapp et al. 2021). These areas are not represented in the climate layers due to landmask constraints in the original HadCM3 simulations, limiting environmental reconstruction in some coastal zones, particularly in northeastern Algeria (Gulf of Bejaia) and northwestern Morocco (Western Rif). Second, Atmosphere-Ocean General Circulation Models (AOGCMs) such as HadCM3 are known to underestimate absolute precipitation values in North Africa, particularly during humid periods (Otto-Bliesner et al., 2017; Valdes et al., 2017). However, we employed clamping during downscaling based on the observed range of modern climatic values, which should mitigate some of these effects. Third, the spatial resolution of the climate model may smooth climatic values across adjacent grid cells, limiting the detection of fine-scale environmental differences between closely spaced sites, such as those in the Rabat–Temara region. While this reduces microregional climatic discrimination, it still captures meaningful variations in broader habitat suitability across MIS 5 substages. Fourth, the climate-envelope models themselves have inherent limitations. They are highly dependent on the distribution of currently known archaeological sites, which remains geographically uneven and comprises a relatively limited number of securely dated MIS 5 occupations. In addition, these models rely exclusively on temperature and precipitation, without accounting for other environmental variables that likely influenced human settlement, particularly the spatial distribution of freshwater resources. The extensive areas of predicted suitability during MIS 5a, despite palaeoclimatic evidence indicating generally drier regional conditions, illustrate these limitations. Together, these results highlight the importance of incorporating reconstructed hydrographic networks, which provide a more realistic representation of landscape accessibility and produce connectivity patterns that are more consistent with both the archaeological record and independent palaeoenvironmental evidence. This integrated approach therefore represents a substantial advance over climate-envelope models based solely on temperature and precipitation. Fifth, the hydrological modelling does not consider the persistence of mega-lakes beyond episodes of peak precipitation or the long-term influence of groundwater reservoirs, both of which may have sustained freshwater availability despite declining rainfall. As a result, the persistence of water bodies may be underestimated in our reconstructions. Sixth, hydrological and optimal path models are sensitive to topographic anomalies, particularly in dune-covered regions where digital elevation models may be less accurate (e.g., Breeze et al., 2015). Nevertheless, the reconstructed stream networks closely match previous hydrographic reconstructions (Drake et al. 2011; Lehner and Grill 2013), reinforcing the robustness of our approach (**SM – Figure 11**). Finally, we acknowledge that optimal paths based on travel time or distance to water cannot capture the full complexity of human movement, which was influenced by many factors beyond landscape features and freshwater availability (e.g., Hewlett 2016; Grove 2009). Nevertheless, the GIS-based framework developed here provides a valuable foundation for exploring human mobility and long-term demographic processes (Bevan and Lake 2013; Howey and Brouwer Burg 2017), and could be extended in future work to incorporate submerged coastal landscapes (Vacchi et al., 2025).

Patterns of precipitation in our model align with previous studies that identify wet phases around ∼125, 105, and 80 ka (**Figure 4**), corresponding to intensifications of the West African Monsoon (e.g, Smith, 2012; Drake et al., 2013; Larrasoaña et al., 2013; Blanchet et al., 2021). Although monsoonal dynamics do not strictly follow MIS substage boundaries, variations in hydrographic extent are clearly visible – particularly in the southern part of the study area – where changes in precipitation had the greatest impact on river activation (**Figure 5; SM – Figure 10**). To our knowledge, this is the first hydrographic reconstruction for MIS 5 to span all North Africa and the first to use palaeoclimatic data to weight the hydrographic reconstruction. Our models reproduce active fluvial systems such as the Sahabi but not the Kufrah river (Coulthard et al. 2013; Paillou et al. 2012) and highlight other Saharan major systems including the Daoura, Soura and Irharhar rivers (**SM – Figure 11**). These north-south or south-north river systems may have connected central Saharan uplands with more humid Mediterranean margins (**Figure 6**), and major streams such as the Draa, Sebou, Moulouya, Chéliff rivers activated during each substages (**Figure 5; SM – Figure 10**). The potential activation of these streams provides context to archaeological evidence of occupation in currently arid areas and is consistent with the widespread distribution of undated surface assemblages located near reconstructed watercourses (**Figures 7** and **9**). This hydrological structure may also help explain patterns of regional population connectivity and fragmentation during MIS 5 (e.g., Scerri et al., 2018).

Lithic assemblages from dated sites (most of which are in Morocco) tend to show a reliance on local raw materials (typically transported <25 km), suggesting localized mobility patterns (Scerri and Spinapolice, 2019). However, exceptional cases such as Adrar Bous (Niger), where exotic materials were transported over distances exceeding 200 km, point to at least some episodes of long-distance movement, potentially along river corridors in southern and central Saharan zones (*ibid*). The timing of these episodes may have been dictated by substage-specific fluctuations in water availability (**Figure 5**), which influenced the extent and distribution of habitable landscapes along the watercourses. The reconstructed hydrological pathways thus provide critical geographical context for understanding the interplay between isolation and interaction in human populations during MIS 5, even though the nature of the interactions between northern, Saharan, and western African regions remains unclear (e.g., Cancellieri et al., 2022). This is especially relevant given emerging archaeological evidence from areas further south (Niang et al., 2023; Ben Arous et al., 2025), which may reshape our understanding of early population mobility and dispersals across this part of the African continent.

Our climate-envelope models reveal spatial shifts in habitat suitability across the five substages of MIS 5 (**Figures 6** and **7**). Vast portions of the Saharan interior were very arid throughout MIS 5, with precipitation often below 100 mm/year (**SM – Figure 10**). Our models indicate that large regions were uninhabited by humans. These results are consistent with field data (Drake et al. 2022) and Holocene data documenting the existence of hyper-arid zones and poor water retention capacity in the eastern Sahara (Kuper and Kröpelin, 2006; Krinner et al., 2012). Northern coastal regions, particularly those benefiting from both Atlantic and Mediterranean influences, appear to have served as macro-refugia (Blinkhorn et al., 2022; Terray et al., 2023), a conclusion supported by our models (**Figures 6** and **24**). Sites along the Atlantic littoral such as Contrebandiers, El Harhoura 2, El Mnasra, and Dar es-Soltan 1-2 exhibit fluctuating conditions between thorn steppe and woodland, while Bizmoune lies near the steppe-woodland threshold. Haua Fteah falls systematically within thorn woodland zones, while climate values reported at Taforalt and Rhafas correspond to more open, thorn steppe environments.

A marked contraction of suitable zones, particularly along the Atlantic littoral, is indicated for substage 5d which coincides with a decline in dated sites. Yet, some locations, such as Contrebandiers, remain repeatedly occupied (**Table 1**), suggesting persistent micro-habitability in this area. MIS 5c and 5b continue this trend of fragmented climate suitability, with small patches of potential habitability mostly concentrated along the circum-Atlantic Mediterranean regions. In contrast, the broad expansion of modelled climate suitability into the central Sahara during MIS 5a appears to be an artefact of the climate-envelope approach. This prediction is largely driven by the inclusion of Uan Afuda, dated to late MIS 5a-early MIS 4 (Boisard et al., 2025), which records the lowest precipitation and highest temperature values in our dataset (**Figure 6**; Appendix). As a result, the MIS 5a climatic envelope extends across much of the Sahara, including regions independently reconstructed as part of the MIS 5e arid belt by Scerri et al. (2014). Rather than reflecting widespread climatic suitability, this discrepancy highlights an important limitation of climate-envelope models based solely on temperature and precipitation. Uan Afuda is in a mountainous setting associated with active streams and drainage basins (**Figure 5**). Although the local temperature and precipitation values approach arid classifications, our hydrological reconstruction indicates the persistent availability of freshwater, which may have sustained favourable local environments. It is therefore plausible that gallery forests or riparian grasslands developed along these drainage systems (Nicoll, 2004; Watrin et al., 2009), consistent with the mosaic model of patchy human occupation proposed by (Garcea, 2004) and Scerri (2017).

As highlighted by Timbrell et al. (2025), MSA occupations in northwest Africa are associated with colder, drier, and less productive environments than those in eastern Africa, with greater temperature seasonality and higher climatic predictability over millennial timescales. These conditions likely encouraged regionally adapted technological strategies which may have been suited to the exploitation of narrow seasonal windows of resource availability. While our climate-envelope models identify zones of persistent climatic suitability based on the habitat preferences of past hunter-gatherers, they do not capture the effects of seasonal variation or hydrological buffering. In this context, our hydrological and mobility models suggest that water availability in semi-arid hinterlands could have facilitated episodic, seasonal or cyclical mobility in otherwise uninhabitable arid zones (**Figures 5** and **7**).

At a broader scale, our modeled path networks reveal potential connectivity paths between the Atlantic littoral (CT) and the Jebel Gharbi region (WG), adding specificity to Scerri’s hypothesis (2013, 2014) of sustained interaction along fluvial corridors (**Figure 7**). In contrast, Cyrenaica appears less integrated with the rest of Northwest Africa, reinforcing a hypothesis of cultural isolation by distance (Scerri et al., 2018). The persistence of distinct cultural traditions in this region may reflect its ecological and geographic particularities (Reade et al., 2016; Boisard et al., 2025). Rather than indicating a uniform pattern of potential connectivity throughout MIS 5, our models provide a framework for evaluating how ecological conditions may have influenced the relative accessibility of locations represented in the archaeological record through time. Zones with dense stream networks and lake potential, especially along the northern coast, may have acted as stable occupation hubs. It is plausible that symbolic behaviors, such as ornament production and pigment use (e.g., Vanhaeren et al., 2006; d’Errico et al., 2009; Barton and d’Errico, 2012) were transmitted along these paths (**Figures 24** and **26**). However, important gaps in our understanding of site distribution patterns in North Africa remain. The archaeological record is still incomplete in several regions, notably Algeria, Tunisia, and submerged coastal margins such as the Gulf of Gabès. Some areas not fully captured by our models – due to the absence of dated sites – nonetheless provide compelling evidence of MSA occupation, including human footprints in northwestern Morocco dating at least to 90 ka (Sedrati et al., 2024) and numerous undated surface assemblages along the Algerian-Tunisian littoral (Boisard and Ben Arous, 2024). These regions, often located near active streams, may also have supported human activity during MIS 5.

## 6. Conclusion

This study adopts a macro-scale approach to investigate how waterscapes and climatic conditions may have structured human occupation and potential mobility in North Africa during MIS 5. By integrating downscaled paleoclimate simulations, hydrological modelling, and GIS-based spatial analyses, we provide a framework for exploring the relationship between environmental change and human movement across the region. Our results suggest that parts of the Central Sahara may have remained intermittently accessible throughout MIS 5 through the reactivation of hydrographic corridors, offering potential pathways linking coastal North Africa with Saharan landscapes. These findings contribute to broader discussions of Late Pleistocene human dispersals within and beyond Africa (Scerri et al., 2018; Beyer et al., 2021) while highlighting the importance of freshwater landscapes in shaping regional accessibility.

Despite limitations in model resolution and the archaeological record, our approach provides a framework for evaluating spatio-temporal changes in landscape accessibility across MIS 5 substages. Future research should focus on refining the chronology of MSA surface assemblages, particularly in Algeria and Tunisia, to better constrain environmental reconstructions and models of human mobility. Undated sites located along reconstructed hydrographic corridors represent promising targets for future investigation, as they may help clarify patterns of occupation and cultural transmission. Finally, while technological similarities are often interpreted as evidence of interaction between groups, they may also arise from recurrent occupations by populations moving through landscapes whose accessibility changed over time. Improved chronological control, combined with technological analyses, will be essential for distinguishing between these alternative scenarios and for further evaluating the role of prehistoric waterscapes in shaping patterns of human occupation and mobility during MIS 5.

## Acknowledgements

SB thanks the University of Colorado – Colorado Springs (UCCS) for their hospitality during the completion of this work.

## Contribution Statement

**Solène Boisard:** Conceptualization, Data curation, Investigation, Methodology, Software, Visualization, Writing – original draft, Writing – review & editing. **Colin D. Wren**: Methodology, Supervision, Resources, Writing – review & editing. **Lucy Timbrell**: Methodology, Software, Writing – review & editing. **Ariane Burke**: Supervision, Writing – review & editing.

## Data availability

All scripts and supplementary material developed for this study are available at: https://doi.org/10.5683/SP3/SED6PH The script used to produce the downscaled climate data is available in Timbrell et al. (2024).

## Appendix

| MIS | sites | level | RawAge | SDev | MIN | MID | MAX | long | lat | N-date | Method | Temp<br>(bio01) | Precip<br>(bio12) |
| --- | --- | --- | --- | --- | --- | --- | --- | --- | --- | --- | --- | --- | --- |
| 5e | Rhafas | S7 | 122500 | 8800 | 114 | 123 | 131 | -1,80 | 34,57 | 1 | OSL | 13,7 | 473 |
| 5e | Ifri n'Ammar | XVII | 130000 | 8000 | 122 | 130 | 138 | -3,28 | 34,69 | 1 | TL | 16,1 | 326 |
| 5e | Dar es-Soltan 1 | Group 1_G1,10 | multiple | multiple | 121 | 128 | 135 | -6,90 | 33,98 | 2 | OSL | 17,2 | 485 |
| 5e | Dar es-Soltan 1 | Group 1_G1,8 | 127540 | 4310 | 123 | 128 | 132 | -6,90 | 33,98 | 1 | OSL | 17,2 | 485 |
| 5e | Dar es-Soltan 2 | 7 | 121700 | 8200 | 114 | 122 | 130 | -6,90 | 33,98 | 1 | OSL | 17,2 | 485 |
| 5e | Contrebandiers | 6c | multiple | multiple | 112 | 126 | 139 | -6,96 | 33,91 | 4 | OSL | 17,4 | 472 |
| 5e | Contrebandiers | 5c | mutuiple | mutiple | 103 | 118 | 133 | -6,96 | 33,91 | 8 | OSL; TL | 17,4 | 472 |
| 5e | Haua Fteah | 832 | multiple | multiple | 116 | 131 | 146 | 22,05 | 32,90 | 2 | IRSL | 18,2 | 373 |
| 5e | Haua Fteah | 827 | 127400 | 8400 | 119 | 128 | 136 | 22,05 | 32,90 | 1 | IRSL | 18,2 | 373 |
| 5e | Haua Fteah | 817 | 126700 | 9100 | 118 | 127 | 136 | 22,05 | 32,90 | 1 | IRSL | 18,2 | 373 |
| 5e | Haua Fteah | 825 | 125500 | 10100 | 115 | 126 | 136 | 22,05 | 32,90 | 1 | IRSL | 18,2 | 373 |
| 5e | Haua Fteah | 828 | 121400 | 7600 | 114 | 122 | 129 | 22,05 | 32,90 | 1 | IRSL | 18,2 | 373 |
| 5e | Haua Fteah | 811 | 119100 | 7200 | 112 | 119 | 126 | 22,05 | 32,90 | 1 | IRSL | 18,2 | 373 |
| 5e | Haua Fteah | 824 | 119100 | 9100 | 110 | 119 | 128 | 22,05 | 32,90 | 1 | IRSL | 18,2 | 373 |
| 5d | Rhafas | 6d | 107000 | 12000 | 95 | 107 | 119 | -1,80 | 34,57 | 1 | OSL | 13,7 | 473 |
| 5d | Dar es-Soltan 1 | Group 2_G2,1 | 113000 | 4010 | 109 | 113 | 117 | -6,90 | 33,98 | 1 | OSL | 17,2 | 485 |
| 5d | Dar es-Soltan 1 | Group 1_G1,7 | 112100 | 8100 | 104 | 112 | 120 | -6,90 | 33,98 | 1 | OSL | 17,2 | 485 |
| 5d | Dar es-Soltan 1 | Group 2_G2,2-6 | 109990 | 4210 | 106 | 110 | 114 | -6,90 | 33,98 | 1 | OSL | 17,2 | 485 |
| 5d | El Harhoura 2 | 9 | 108100 | 5900 | 102 | 108 | 114 | -6,93 | 33,95 | 1 | OSL | 17,3 | 475 |
| 5d | El Mnasra | 9c | multiple | multiple | 102 | 109 | 115 | -6,95 | 33,93 | 2 | OSL | 17,3 | 475 |
| 5d | Contrebandiers | V-1b | 113000 | 7000 | 106 | 113 | 120 | -6,96 | 33,91 | 1 | OSL | 17,4 | 472 |
| 5d | Contrebandiers | 4d | multiple | multiple | 90 | 108 | 126 | -6,96 | 33,91 | 4 | OSL | 17,4 | 472 |
| 5d | Contrebandiers | V-2 | 107000 | 9000 | 98 | 107 | 116 | -6,96 | 33,91 | 1 | OSL | 17,4 | 472 |
| 5d | Contrebandiers | 5 | multiple | multiple | 97 | 106 | 116 | -6,96 | 33,91 | 2 | OSL | 17,4 | 472 |
| 5d | Contrebandiers | 4c | multiple | multiple | 96 | 105 | 114 | -6,96 | 33,91 | 2 | OSL | 17,4 | 472 |
| 5c | Rhafas | 3b | multiple | multiple | 66 | 92 | 118 | -1,80 | 34,57 | 5 | OSL; TL | 13,7 | 473 |
| 5c | Taforalt | S1-S6-CalcGr | multiple | multiple | 84 | 100 | 115 | -2,40 | 34,81 | 10 | U-Th | 15,1 | 353 |
| 5c | Taforalt | Group F | multiple | multiple | 77 | 95 | 113 | -2,40 | 34,81 | 2 | OSL | 15,1 | 353 |
| 5c | Dar es-Soltan 2 | 5 | 101400 | 11100 | 90 | 102 | 113 | -6,90 | 33,98 | 1 | OSL | 17,2 | 485 |
| 5c | El Harhoura 2 | 8 | 92000 | 11000 | 81 | 92 | 103 | -6,93 | 33,95 | 1 | US-ESR | 17,3 | 475 |
| 5c | El Mnasra | 7 | multiple | multiple | 70 | 100 | 130 | -6,95 | 33,93 | 7 | OSL | 17,3 | 475 |
| 5c | El Mnasra | 8b | multiple | multiple | 69 | 94 | 119 | -6,95 | 33,93 | 6 | OSL; US-ESR | 17,3 | 475 |
| 5c | El Mnasra | 8c-d | multiple | multiple | 68 | 92 | 115 | -6,95 | 33,93 | 4 | OSL; US-ESR | 17,3 | 475 |
| 5c | Contrebandiers | IV-2b | 101000 | 9000 | 92 | 101 | 110 | -6,96 | 33,91 | 1 | OSL | 17,4 | 472 |
| 5c | Contrebandiers | 4 d-e | multiple | multiple | 89 | 100 | 110 | -6,96 | 33,91 | 2 | OSL | 17,4 | 472 |
| 5c | Contrebandiers | 5a | multiple | multiple | 73 | 100 | 127 | -6,96 | 33,91 | 2 | OSL; TL | 17,4 | 472 |
| 5c | Contrebandiers | 5b | multiple | multiple | 75 | 100 | 124 | -6,96 | 33,91 | 2 | OSL; TL | 17,4 | 472 |
| 5c | Contrebandiers | IV-2a | 96000 | 8000 | 88 | 96 | 104 | -6,96 | 33,91 | 1 | OSL | 17,4 | 472 |
| 5c | Contrebandiers | 4 | 92000 | 6000 | 86 | 92 | 98 | -6,96 | 33,91 | 1 | OSL | 17,4 | 472 |
| 5c | Bizmoune | 4a | 102980 | 9620 | 93 | 103 | 113 | -9,56 | 31,67 | 1 | U-Th | 17,8 | 315 |
| 5c | Bizmoune | 3b-3inf | multiple | multiple | 76 | 94 | 111 | -9,56 | 31,67 | 2 | OSL; U-Th | 17,8 | 315 |
| 5c | Haua Fteah | 770 | 103800 | 7400 | 96 | 104 | 111 | 22,05 | 32,90 | 1 | IRSL | 18,2 | 373 |
| 5c | Haua Fteah | 798 | 102200 | 8700 | 94 | 103 | 111 | 22,05 | 32,90 | 1 | IRSL | 18,2 | 373 |
| 5b | Rhafas | 3a | multiple | multiple | 63 | 91 | 118 | -1,80 | 34,57 | 4 | OSL; TL | 13,7 | 473 |
| 5b | Rhafas | S6 | 86400 | 4900 | 81 | 86 | 91 | -1,80 | 34,57 | 1 | OSL | 13,7 | 473 |
| 5b | Dar es-Soltan 1 | Group 3_G3,1 | 88520 | 3430 | 85 | 89 | 92 | -6,90 | 33,98 | 1 | OSL | 17,2 | 485 |
| 5b | Dar es-Soltan 1 | Group 3_G3,3 | 85960 | 3290 | 82 | 86 | 89 | -6,90 | 33,98 | 1 | OSL | 17,2 | 485 |
| 5b | El Mnasra | 9 | 90000 | 5000 | 85 | 90 | 95 | -6,95 | 33,93 | 1 | US-ESR | 17,3 | 475 |
| 5b | El Mnasra | 8a | multiple | multiple | 62 | 90 | 117 | -6,95 | 33,93 | 4 | OSL; US-ESR | 17,3 | 475 |
| 5b | Contrebandiers | 3a | multiple | multiple | 73 | 90 | 106 | -6,96 | 33,91 | 3 | OSL; TL | 17,4 | 472 |
| 5b | Contrebandiers | IV-2 | multiple | multiple | 69 | 87 | 104 | -6,96 | 33,91 | 6 | OSL; TL | 17,4 | 472 |
| 5b | Haua Fteah | 776 | 89500 | 5300 | 84 | 90 | 95 | 22,05 | 32,90 | 1 | OSL | 18,2 | 373 |
| 5a | Rhafas | 3 | 82000 | 8000 | 74 | 82 | 90 | -1,80 | 34,57 | 1 | TL | 13,7 | 473 |
| 5a | Taforalt | S1-S6-GroupE | multiple | multiple | 68 | 79 | 89 | -2,40 | 34,81 | 2 | OSL | 15,1 | 353 |
| 5a | Dar es-Soltan 1 | Group 3_G3,4 | 82790 | 3310 | 79 | 83 | 86 | -6,90 | 33,98 | 1 | OSL | 17,2 | 485 |
| 5a | Dar es-Soltan 1 | Group 3_G3,11 | 77150 | 2960 | 74 | 77 | 80 | -6,90 | 33,98 | 1 | OSL | 17,2 | 485 |
| 5a | Bizmoune | 3 | multiple | multiple | 64 | 72 | 80 | -9,56 | 31,67 | 7 | OSL | 17,8 | 315 |
| 5a | Haua Fteah | 528 | 72800 | 5200 | 67 | 73 | 78 | 22,05 | 32,90 | 1 | IRSL | 18,2 | 373 |
| 5a | Haua Fteah | 525/565 | 71900 | 7600 | 64 | 72 | 80 | 22,05 | 32,90 | 1 | OSL | 18,2 | 373 |
| 5a | Haua Fteah | 524 | 71200 | 5800 | 65 | 71 | 77 | 22,05 | 32,90 | 1 | IRSL | 18,2 | 373 |
| 5a | Uan Afuda | dune | multiple | multiple | 62 | 73 | 83 | 10,50 | 24,87 | 3 | OSL; TL | 23,9 | 19 |

## References

Barton, N., Errico, F. d’, 2012. North Africans origins of symbolically mediated behaviour and the Aterian. In: Elias, S. (Ed.), Origins of Human Innovation and Creativity, Developments in Quaternary Science. Elsevier B.B., pp. 23–34.

Ben Arous, E., Blinkhorn, J.A., Elliott, S., Kiahtipes, C.A., N’zi, C.D., Bateman, M.D., Duval, M., Roberts, P., Patalano, R., Blackwood, A.F., Niang, K., Kouamé, E.A., Lebato, E., Hallett, E., Cerasoni, J.N., Scott, E., Ilgner, J., Alonso Escarza, M.J., Guédé, F.Y., Scerri, E.M.L., 2025. Humans in Africa’s wet tropical forests 150 thousand years ago. Nature 1–6. doi:10.1038/s41586-025-08613-y

Bevan, A., Lake, M. (Eds.), 2013. Computational approaches to archaeological spaces, Publications of the Institute of Archaeology, University College London. Left Coast Press, Walnut Creek, California.

Beyer, R.M., Krapp, M., Eriksson, A., Manica, A., 2021. Climatic windows for human migration out of Africa in the past 300,000 years. Nature Communications 12, 4889. doi:10.1038/s41467-021-24779-1

Blanchet, C.L., Osborne, A.H., Tjallingii, R., Ehrmann, W., Friedrich, T., Timmermann, A., Brückmann, W., Frank, M., 2021. Drivers of river reactivation in North Africa during the last glacial cycle. Nature Geoscience 14, 97–103. doi:10.1038/s41561-020-00671-3

Blinkhorn, J., Timbrell, L., Grove, M., Scerri, E.M.L., 2022. Evaluating refugia in recent human evolution in Africa. Philosophical Transactions of the Royal Society B: Biological Sciences 377, 20200485. doi:10.1098/rstb.2020.0485

Blome, M.W., Cohen, A.S., Tryon, C.A., Brooks, A.S., Russell, J., 2012. The environmental context for the origins of modern human diversity: a synthesis of regional variability in African climate 150,000-30,000 years ago. Journal of Human Evolution 62, 563–592. doi:10.1016/j.jhevol.2012.01.011

Boisard, S., Ben Arous, E., 2024. A Critical Inventory and Associated Chronology of the Middle Stone Age and Later Stone Age in Northwest Africa. Journal of Open Archaeology Data 12, 5. doi:10.5334/joad.121

Boisard, S., Wren, C.D., Timbrell, L., Burke, A., 2025. Climate frameworks for the Middle Stone Age and Later Stone Age in Northwest Africa. Quaternary International 716, 109593. doi:10.1016/j.quaint.2024.109593

Breeze, P.S., Drake, N.A., Groucutt, H.S., Parton, A., Jennings, R.P., White, T.S., Clark-Balzan, L., Shipton, C., Scerri, E.M.L., Stimpson, C.M., Crassard, R., Hilbert, Y., Alsharekh, A., Al-Omari, A., Petraglia, M.D., 2015. Remote sensing and GIS techniques for reconstructing Arabian palaeohydrology and identifying archaeological sites. Quaternary International, Green Arabia: Human Prehistory at the Cross-roads of Continents 382, 98–119. doi:10.1016/j.quaint.2015.01.022

Cancellieri, E., 2021. A tentative tale of Stone Age human dynamics in Pleistocene south-western Libya (central Sahara). Libyan Studies 52, 36–53. doi:10.1017/lis.2021.18

Cancellieri, E., Bel Hadj Brahim, H., Ben Nasr, J., Ben Fraj, T., Boussoffara, R., Di Matteo, M., Mercier, N., Marnaoui, M., Monaco, A., Richard, M., Mariani, G.S., Scancarello, O., Zerboni, A., Lernia, S. di, 2022. A late Middle Pleistocene Middle Stone Age sequence identified at Wadi Lazalim in southern Tunisia. Scientific Reports 12, 3996. doi:10.1038/s41598-022-07816-x

Coulthard, T.J., Ramirez, J.A., Barton, N., Rogerson, M., Brücher, T., 2013. Were Rivers Flowing across the Sahara During the Last Interglacial? Implications for Human Migration through Africa. PLOS ONE 8, e74834. doi:10.1371/journal.pone.0074834

Creveling, J.R., Mitrovica, J.X., Clark, P.U., Waelbroeck, C., Pico, T., 2017. Predicted bounds on peak global mean sea level during marine isotope stages 5a and 5c. Quaternary Science Reviews 163, 193–208. doi:10.1016/j.quascirev.2017.03.003

Drake, N., Blench, R.M., Armitage, S.J., Bristow, C.S., White, K.H., 2011. Ancient watercourses and biogeography of the Sahara explain the peopling of the desert. Proceedings of the National Academy of Science 108, 458–462. doi:www.pnas.org/cgi/doi/10.1073/pnas.1012231108

Drake, N., Breeze, P., Parker, A., 2013. Palaeoclimate in the Saharan and Arabian deserts during the Middle Palaeolithic and the potential for hominin dispersals. Quaternary International 300, 48–61.

Drake, N.A., Candy, I., Breeze, P., Armitage, S.J., Gasmi, N., Schwenninger, J.L., Peat, D., Manning, K., 2022. Sedimentary and geomorphic evidence of Saharan megalakes: A synthesis. Quaternary Science Reviews 276, 107318. doi:10.1016/j.quascirev.2021.107318

Errico, F. d’, Vanhaeren, M., Barton, N., Bouzouggar, A., Mienis, H., Richter, D., Hublin, J.-J., McPherron, S.P., Lozouet, P., 2009a. Additional evidence on the use of personal ornaments in the Middle Paleolithic of North Africa. Proceedings of the National Academy of Science 106, 16051–6.

Errico, F. d’, Vanhaeren, M., Barton, N., Bouzouggar, A., Mienis, H., Richter, D., Hublin, J.-J., McPherron, S.P., Lozouet, P., 2009b. Additional evidence on the use of personal ornaments in the Middle Paleolithic of North Africa. Proceedings of the National Academy of Science 106, 16051–6.

Fick, S., Hijmans, R., 2017. WorldClim 2: New 1-km spatial resolution climate surfaces for global land areas. International Journal of Climatology 37. doi:10.1002/joc.5086

Garcea, E.A.A., 2004. Crossing Deserts and Avoiding Seas: Aterian North African-European Relations. Journal of Anthropological Research 60, 27–53.

Gasse, F., 2000. Hydrological changes in the African tropics since the Last Glacial Maximum. Quaternary Science Reviews 19, 189–211. doi:10.1016/S0277-3791(99)00061-X

Grove, M., 2009. Hunter–gatherer movement patterns: Causes and constraints. Journal of Anthropological Archaeology 28, 222–233. doi:10.1016/j.jaa.2009.01.003

Herzog, I., 2013. The Potential and Limits of Optimal Path Analysis. In: Computational Approaches to Archaeological Spaces. Routledge.

Hewlett, B.S., 2016. Social learning and Innovation in Hunter-Gatherers. In: Terashima, H., Hewlett, B.S. (Eds.), Social Learning and Innovation in Contemporary Hunter-Gatherers: Evolutionary and Ethnographic Perspectives, Replacement of Neanderthals by Modern Humans Series. Springer Japan, Tokyo, pp. 1–15. doi:10.1007/978-4-431-55997-9_17

Holdridge, L.R., 1967. Life Zone Ecology. Tropical Science Center.

Howey, M.C.L., Brouwer Burg, M., 2017. Assessing the state of archaeological GIS research: Unbinding analyses of past landscapes. Journal of Archaeological Science, Archaeological GIS Today: Persistent Challenges, Pushing Old Boundaries, and Exploring New Horizons 84, 1–9. doi:10.1016/j.jas.2017.05.002

Hublin, J.-J., McPherron, S.P. (Eds.), 2012. Modern Origins: A North African Perspective, New-York: Springer. ed.

Krapp, M., Beyer, R.M., Edmundson, S.L., Valdes, P.J., Manica, A., 2021. A statistics-based reconstruction of high-resolution global terrestrial climate for the last 800,000 years. Scientific Data 8, 228. doi:10.1038/s41597-021-01009-3

Krinner, G., Lézine, A.-M., Braconnot, P., Sepulchre, P., Ramstein, G., Grenier, C., Gouttevin, I., 2012. A reassessment of lake and wetland feedbacks on the North African Holocene climate. Geophysical Research Letters 39. doi:10.1029/2012GL050992

Kukla, G.J., Bender, M.L., Beaulieu, J.-L. de, Bond, G., Broecker, W.S., Cleveringa, P., Gavin, J.E., Herbert, T.D., Imbrie, J., Jouzel, J., Keigwin, L.D., Knudsen, K.-L., McManus, J.F., Merkt, J., Muhs, D.R., Müller, H., Poore, R.Z., Porter, S.C., Seret, G., Shackleton, N.J., Turner, C., Tzedakis, P.C., Winograd, I.J., 2002. Last Interglacial Climates. Quaternary Research 58, 2– 13. doi:10.1006/qres.2001.2316

Kuper, R., Kröpelin, S., 2006. Climate-Controlled Holocene Occupation in the Sahara: Motor of Africa’s Evolution. Science 313, 803–807.

Larrasoaña, J.C., Roberts, A.P., Rohling, E.J., 2013. Dynamics of Green Sahara Periods and Their Role in Hominin Evolution. PLOS ONE 8, e76514. doi:10.1371/journal.pone.0076514

Lehner, B., Grill, G., 2013. Global river hydrography and network routing: baseline data and new approaches to study the world’s large river systems. Hydrological Processes 27, 2171– 2186. doi:10.1002/hyp.9740

Niang, K., Blinkhorn, J., Bateman, M.D., Kiahtipes, C.A., 2023. Longstanding behavioural stability in West Africa extends to the Middle Pleistocene at Bargny, coastal Senegal. Nature Ecology & Evolution 1–11. doi:10.1038/s41559-023-02046-4

Nicoll, K., 2004. Recent environmental change and prehistoric human activity in Egypt and Northern Sudan. Quaternary Science Reviews 23, 561–580. doi:10.1016/j.quascirev.2003.10.004

NOAA National Centers for Environmental Information, 2022. ETOPO 2022 15 Arc-Second Global Relief Model. 10.25921/fd45-gt74

O’Mara, N., Skonieczny, C., McGee, D., Winckler, G., Bory, A., Bradtmiller, L., Malaizé, B., Polissar, P., 2022. Pleistocene drivers of Northwest African hydroclimate and vegetation. Nature Communications 13. doi:10.1038/s41467-022-31120-x

Osborne, A.H., Vance, D., Rohling, E.J., Barton, N., Rogerson, M., Fello, N., 2008. A humid corridor across the Sahara for the migration of early modern humans out of Africa 120,000 years ago. Proceedings of the National Academy of Science 105, 16444–16447.

Otto-Bliesner, B.L., Braconnot, P., Harrison, S.P., Lunt, D.J., Abe-Ouchi, A., Albani, S., Bartlein, P.J., Capron, E., Carlson, A.E., Dutton, A., Fischer, H., Goelzer, H., Govin, A., Haywood, A., Joos, F., LeGrande, A.N., Lipscomb, W.H., Lohmann, G., Mahowald, N., Nehrbass-Ahles, C., Pausata, F.S.R., Peterschmitt, J.-Y., Phipps, S.J., Renssen, H., Zhang, Q., 2017. The PMIP4 contribution to CMIP6 – Part 2: Two interglacials, scientific objective and experimental design for Holocene and Last Interglacial simulations. Geoscientific Model Development 10, 3979–4003. doi:10.5194/gmd-10-3979-2017

Pausata, F.S.R., Gaetani, M., Messori, G., Berg, A., Maia de Souza, D., Sage, R.F., deMenocal, P.B., 2020. The Greening of the Sahara: Past Changes and Future Implications. One Earth 2, 235–250. doi:10.1016/j.oneear.2020.03.002

R Studio Team, 2020. RStudio: Integrated Development for R. RStudio.

Railsback, L.B., Gibbard, P.L., Head, M.J., Voarintsoa, N.R.G., Toucanne, S., 2015. An optimized scheme of lettered marine isotope substages for the last 1.0 million years, and the climatostratigraphic nature of isotope stages and substages. Quaternary Science Reviews 111, 94–106. doi:10.1016/j.quascirev.2015.01.012

Reade, H., Barker, G., Stevens, R.E., O’Connell, T.C., 2016. Pleistocene and Holocene palaeoclimates in the Gebel Akhdar (Libya) estimated using herbivore tooth enamel oxygen isotope compositions. Quaternary International.

Richter, D., Grün, R., Joannes-Boyau, R., Steele, T.E., Amani, F., Rué, M., Fernandes, P., Raynal, J.-P., Geraads, D., Ben-Ncer, A., Hublin, J.-J., McPherron, S.P., 2017. The age of the hominin fossils from Jebel Irhoud, Morocco, and the origins of the Middle Stone Age. Nature 546, 293–296. doi:10.1038/nature22335

Scerri, E.M.L., 2017. The North African Middle Stone Age and its place in recent human evolution. Evolutionary Anthropology: Issues, News, and Reviews 26, 119–135. doi:10.1002/evan.21527

Scerri, E.M.L., Spinapolice, E.E., 2019. Lithics of the North African Middle Stone Age: assumptions, evidence and future directions. Journal of Anthropological Sciences 1–36. doi:10.4436/JASS.97002

Scerri, E.M.L., Will, M., 2023. The revolution that still isn’t: The origins of behavioral complexity in Homo sapiens. Journal of Human Evolution 179, 103358. doi:10.1016/j.jhevol.2023.103358

Scerri, E.M.L., Drake, N., Jennings, R.P., Groucutt, H.S., 2014. Earliest evidence for the structure of Homo sapiens populations in Africa. Quaternary Science Reviews 101, 207–216.

Scerri, E.M.L., Thomas, M.G., Manica, A., Gunz, P., Stock, J.T., Stringer, C., Grove, M., Groucutt, H.S., Timmermann, A., Rightmire, G.P., Errico, F. d’, Tryon, C.A., Drake, N.A., Brooks, A.S., Dennell, R.W., Durbin, R., Henn, B.M., Lee-Thorp, J., deMenocal, P., Petraglia, M.D., Thompson, J.C., Scally, A., Chikhi, L., 2018a. Did Our Species Evolve in Subdivided Populations across Africa, and Why Does It Matter? Trends in Ecology & Evolution 33, 582–594. doi:10.1016/j.tree.2018.05.005

Scerri, E.M.L., Thomas, M.G., Manica, A., Gunz, P., Stock, J.T., Stringer, C., Grove, M., Groucutt, H.S., Timmermann, A., Rightmire, G.P., Errico, F. d’, Tryon, C.A., Drake, N.A., Brooks, A.S., Dennell, R.W., Durbin, R., Henn, B.M., Lee-Thorp, J., deMenocal, P., Petraglia, M.D., Thompson, J.C., Scally, A., Chikhi, L., 2018b. Did Our Species Evolve in Subdivided Populations across Africa, and Why Does It Matter? 33, 582–594. doi:10.1016/j.tree.2018.05.005

Scerri, E.M.L., Chikhi, L., Thomas, M.G., 2019. Beyond multiregional and simple out-of-Africa models of human evolution. Nature Ecology & Evolution 3, 1370–1372. doi:10.1038/s41559-019-0992-1

Sedrati, M., Morales, J.A., Duveau, J., M’rini, A.E., Mayoral, E., Díaz-Martínez, I., Anthony, E.J., Bulot, G., Sedrati, A., Le Gall, R., Santos, A., Rivera-Silva, J., 2024. A Late Pleistocene hominin footprint site on the North African coast of Morocco. Scientific Reports 14, 1962. doi:10.1038/s41598-024-52344-5

Smith, J.R., 2012. Spatial and Temporal Variation in the Nature of Pleistocene Pluvial Phase Environments Across North Africa. In: Hublin, J.-J., McPherron, S.P. (Eds.), Modern Origins: A North African Perspective, Vertebrate Paleobiology and Paleoanthropology. Springer Netherlands, Dordrecht, pp. 35–47. doi:10.1007/978-94-007-2929-2_3

Stewart, J.R., Stringer, C.B., 2012. Human Evolution Out of Africa: The Role of Refugia and Climate Change. Science 335, 1317–1321. doi:10.1126/science.1215627

Terray, L., Stoetzel, E., Ben Arous, E., Kageyama, M., Cornette, R., Braconnot, P., 2023. Refinement of the environmental and chronological context of the archeological site El Harhoura 2 (Rabat, Morocco) using paleoclimatic simulations. Climate of the Past 19, 1245–1263. doi:10.5194/cp-19-1245-2023

Timbrell, L., Blinkhorn, J., Colucci, M., Leonardi, M., Chevalier, M., Grove, M., Scerri, E., Manica, A., 2024. More is not always better: downscaling climate model outputs from 30 to 5-minute resolution has minimal impact on coherence with Late Quaternary proxies. Climate of the Past Discussions 1–21. doi:10.5194/cp-2024-53

Timmermann, A., Friedrich, T., 2016. Late Pleistocene climate drivers of early human migration. Nature 538, 92–95. doi:10.1038/nature19365

Tobler, W., 1993. Three Presentations on Geographical Analysis and Modeling: Non-Isotropic Geographic Modeling; Speculations on the Geometry of Geography; and Global Spatial Analysis (93–1).

USGS, 2017. Shuttle Radar Topography Mission (SRTM) 1 Arc-Second Global. doi:10.5066/F7PR7TFT

Vacchi, M., Shaw, T., Anthony, E., Spada, G., Melini, D., Li, T., Cahill, N., Horton, B., 2025. Sea level since the Last Glacial Maximum from the Atlantic coast of Africa. Nature Communications 16. doi:10.1038/s41467-025-56721-0

Valdes, P.J., Armstrong, E., Badger, M.P.S., Bradshaw, C.D., Bragg, F., Crucifix, M., Davies-Barnard, T., Day, J.J., Farnsworth, A., Gordon, C., Hopcroft, P.O., Kennedy, A.T., Lord, N.S., Lunt, D.J., Marzocchi, A., Parry, L.M., Pope, V., Roberts, W.H.G., Stone, E.J., Tourte, G.J.L., Williams, J.H.T., 2017. The BRIDGE HadCM3 family of climate models: HadCM3@Bristol v1.0. Geoscientific Model Development 10, 3715–3743. doi:10.5194/gmd-10-3715-2017

Van Peer, P., 2016. Technological Systems, Population Dynamics, and Historical Process in the MSA of Northern Africa. In: Jones, S.C., Stewart, B.A. (Eds.), Africa from MIS 6-2: Population Dynamics and Paleoenvironments, Vertebrate Paleobiology and Paleoanthropology. Springer Netherlands, Dordrecht, pp. 147–159. doi:10.1007/978-94-017-7520-5_8

Vanhaeren, M., Errico, F. d’, Stringer, C., James, S.L., Todd, J.A., Mienis, H.K., 2006. Middle Paleolithic Shell Beads in Israel and Algeria. Science 312, 1785–1788. doi:10.1126/science.1128139

Vidal, C.M., Lane, C.S., Asrat, A., Barfod, D.N., Mark, D.F., Tomlinson, E.L., Tadesse, A.Z., Yirgu, G., Deino, A., Hutchison, W., Mounier, A., Oppenheimer, C., 2022. Age of the oldest known Homo sapiens from eastern Africa. Nature 601, 579–583. doi:10.1038/s41586-021-04275-8

Watrin, J., Lézine, A.-M., Hély, C., 2009. Plant migration and plant communities at the time of the “green Sahara.” Comptes Rendus Geoscience, Histoire climatique des déserts d’Afrique et d’Arabie 341, 656–670. doi:10.1016/j.crte.2009.06.007

