## Supplementary Material for "Reconstructing Prehistoric Waterscapes and Human Mobility during MIS 5 in North Africa"

**Supplementary Material** (also available at https://doi.org/10.5683/SP3/SED6PH)

***Lakes***

MIS 5 lakes reconstruction differ regarding lake extent. The three largest megalakes considered active during MIS 5 include: Chotts Megalake system (Tunisia-Algeria): ~30 000 km², consisting of three interconnected basins (Drake et al., 2022); Ahnet-Mouydir Megalake (southern Algeria): at minimum ~50 000 km², U-Th dated to 102 ± 16–14 ka (Kadda; Causse et al., 1989) and 92 ± 20–18 ka (Azzel Matti; Causse et al., 1988), though its exact extent remains uncertain (Drake et al., 2022) and Mega Fezzan Lake (western Libya) which reached an estimated ~1,600 km² at its maximum extent (Armitage et al., 2007; Drake et al., 2018, 2022). In addition to these large lakes, this study incorporates all smaller lakes included in Drake et al.'s (2011) reconstruction. These smaller lakes are assumed to have been active during MIS 5, based on their presence during the Holocene - a period generally considered less humid than MIS 5 (Drake et al. 2022).

***Optimal Path Analysis***

Four different scenarios were tested to assess how different cost variables influenced movement pathways by varying the weights assigned to each factor (see **Table 3**).

***Slope***

The slope layer used in the optimal path analysis was derived from a Digital Elevation Model (DEM) and processed to represent travel difficulty based on terrain steepness. To model the impact of slope on movement, we applied Tobler's Hiking Function, which estimates travel speed (km/h) as a function of slope. Since optimal path modeling requires a cost surface where higher values indicate greater difficulty, we converted travel speed into travel cost, assigning lower values to flatter terrain and higher values to steeper slopes. First, we computed the slope layer (originally in degrees) in radians. Then, we applied Tobler's function to the slope raster using Raster Calculator, generating a travel speed raster, where higher values represent faster movement on flatter terrain:

6 * Exp(-3.5 * Abs(Tan("slope_radians") + 0.05))

We then inverted the speed raster to create a travel cost raster, where lower values correspond to easier movement (flat terrain) and higher values indicate more difficult movement (steep slopes):

1 / (6 * Exp(-3.5 * Abs(Tan("slope_radians") + 0.05)))

The final slope cost layer (tobler_cost) was then integrated into the Weighted Sum function alongside the water distance and precipitation layers to compute the optimal path analysis.

***Precipitation***

We calculated in R the mean total annual precipitation for each MIS 5 substage using paleoclimate datasets (**Figure 10**). Given the physiological constraints of the human body (Wolf et al., 2022), extremely dry areas were expected to be less favorable for movement. To account for this, we applied a linear transformation that assigns higher values to drier regions and lower values to wetter areas, using the minimum and maximum precipitation values from the dataset of each substage – example:

100 - (("mean_precip_5a" - 2.15419) / (1376.54 - 2.15419) * 99)

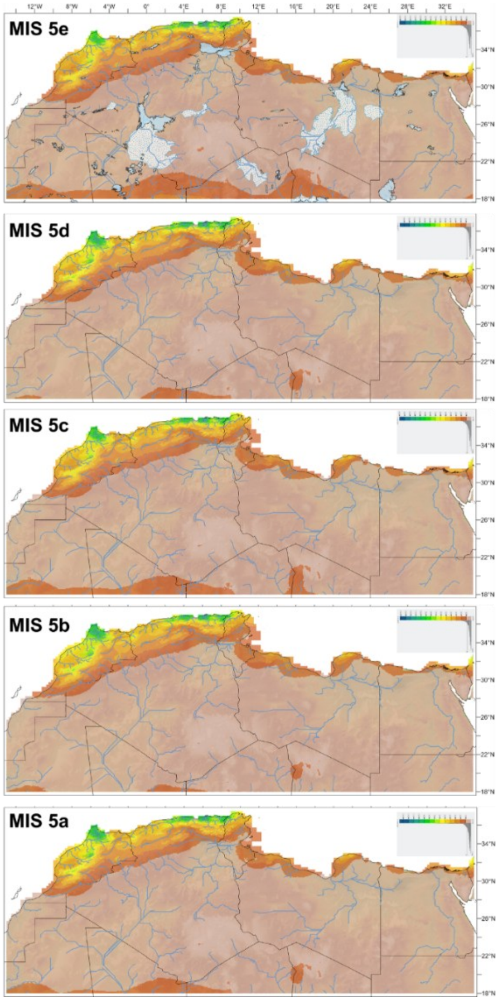


**Figure 10.** Comparison of the means of total annual precipitation and hydrographic modeling between substages (flow accumulation 10,000,000).

***Water distance***

For each hydrological reconstruction corresponding to a specific MIS 5 substage, we generated a distance accumulation raster to quantify proximity to water sources. The hydrological network was used as a direct input feature in the Distance Accumulation tool in ArcGIS Pro, allowing us to compute a raster where: low values indicate areas near water and higher values represent greater distances from rivers.

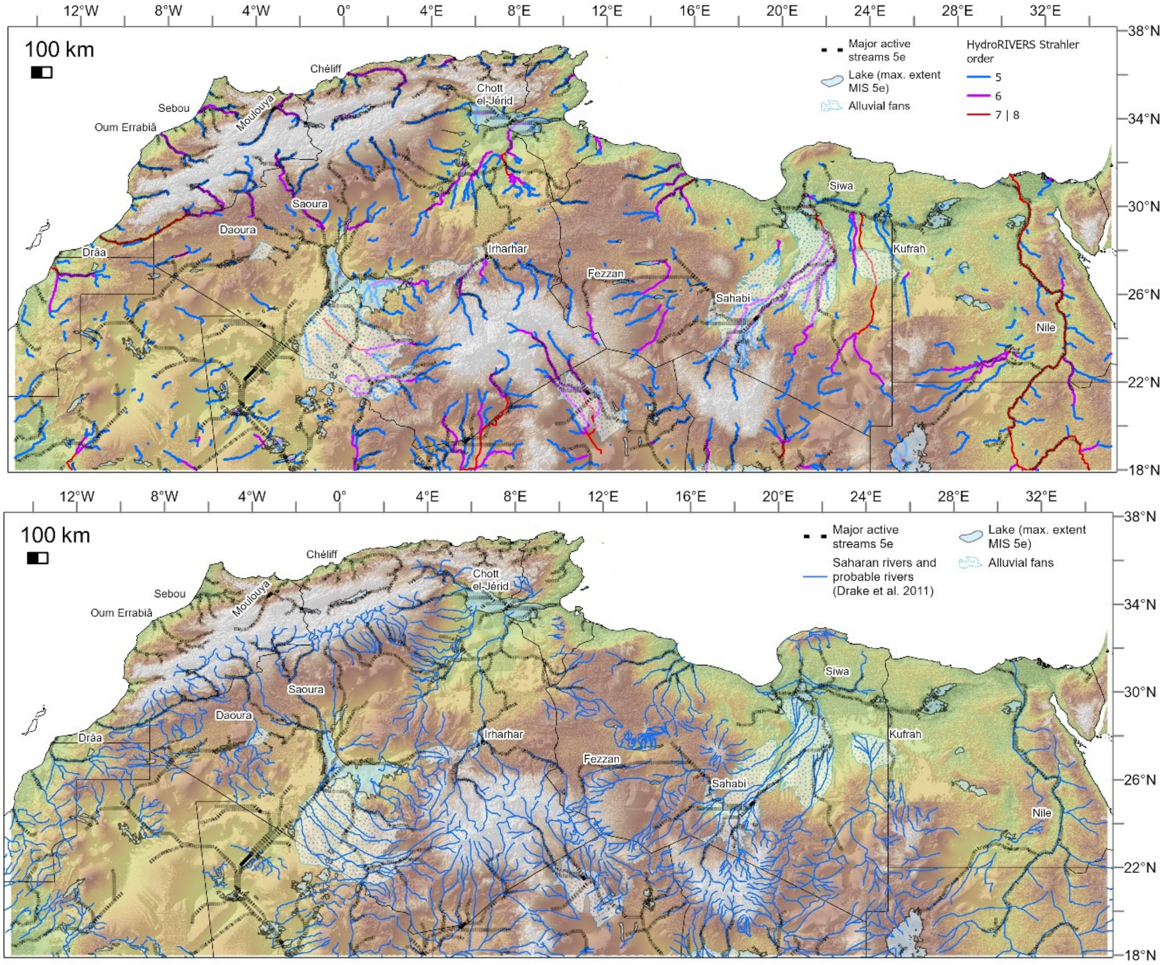


**Figure 11.** Comparison of MIS 5e hydrographic reconstruction with Drake et al. (2011) and HydroRIVERS reconstruction with selection of high Strahler orders (Lehner and Grill, 2013).

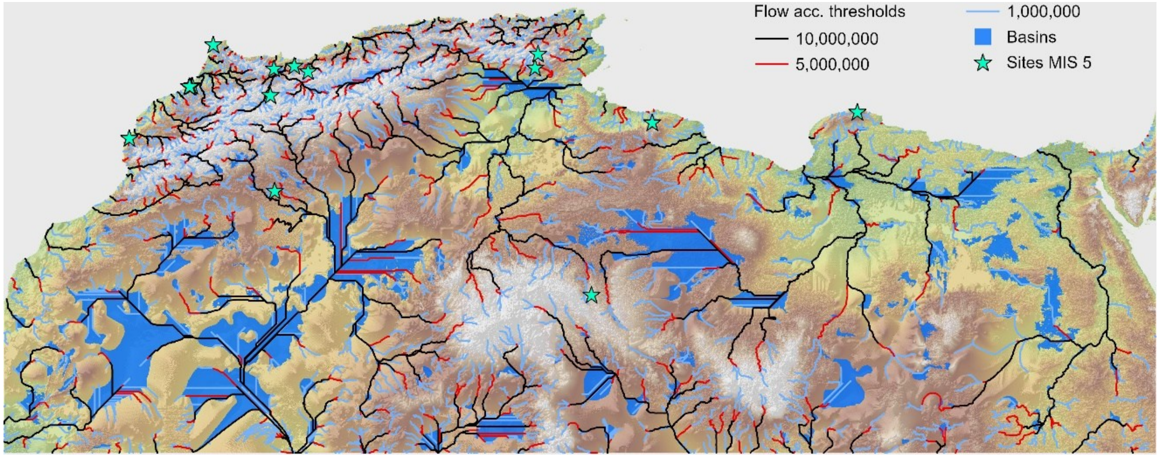


**Figure 12.** Comparison of distinct flow accumulation threshold for MIS 5e hydrographic reconstruction.

### Data Normalization

To combine the three environmental layers (slope, water distance, and precipitation) into a single weighted cost surface, all layers were normalized on a scale from 1 to 100:

<u>Slope</u>: 1 represents flat terrain, 100 represents the steepest slopes.
<u>Precipitation</u>: 1 represents the most humid regions, 100 represents the driest areas.
<u>Water distance</u>: 1 represents areas closest to water, 100 represents the farthest areas.

The final weight raster produced a composite cost layer, where lower values (1) indicate the most favorable areas for movement, and higher values (100) represent the least favorable areas.

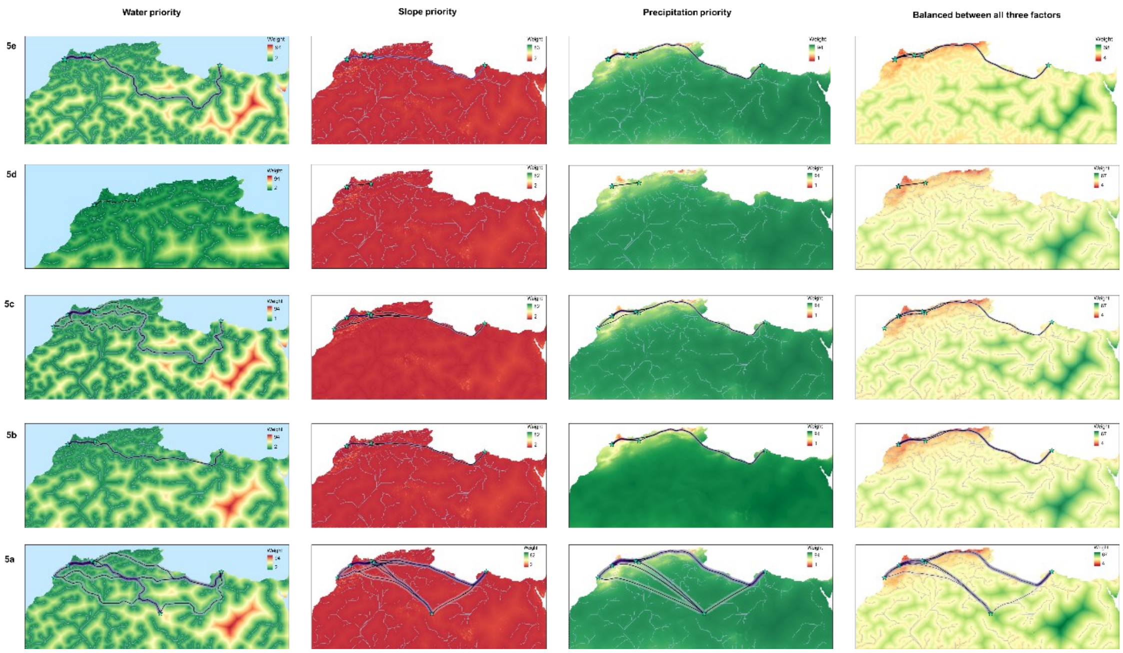


**Figure 13.** Maps display weighted layers and optimal paths according to different environmental priorities.
